# Biased connectivity of brain-wide inputs to ventral subiculum output neurons

**DOI:** 10.1101/726406

**Authors:** RWS Wee, AF MacAskill

## Abstract

The ventral subiculum (vS) of the mouse hippocampus coordinates diverse behaviours through heterogeneous populations of projection neurons. These neurons transmit signals to multiple brain regions by integrating thousands of local and long-range synaptic inputs. However, whether each population is selectively innervated by different afferent input remains unknown. To address this question, we employed projection-specific rabies tracing to study the input-output relationship of vS output neurons. Analysis of brain-wide inputs reveals quantitative input differences that can be explained by the spatial location of postsynaptic neurons along the proximal-distal axis of vS and the identity of the downstream target. Further, the input from nucleus reuniens, an area thought to underlie vS and prefrontal cortex (PFC) reciprocal connectivity, is unexpectedly biased away from PFC-projecting vS neurons. Overall, we reveal prominent heterogeneity in brain-wide inputs to the vS parallel output circuitry, providing a basis for the selective control of individual projections during behaviour.

## Introduction

The rodent hippocampus coordinates a wide array of behaviours (Strange et al., 2014), from spatial navigation (O’Keefe and Dostrovsky, 1971) to decision-making under approach-avoidance conflict (Ciocchi et al., 2015; Jimenez et al., 2018) and reward processing (Ciocchi et al., 2015). A central hypothesis of how the hippocampus might participate in such diverse behaviours is the presence of heterogeneous principal neurons that differ widely in their gene expression (Cembrowski et al., 2016; Strange et al., 2014), electrophysiological properties (Kim and Spruston, 2011) and behavioural function (Cembrowski et al., 2018a; 2018b; Ciocchi et al., 2015; Strange et al., 2014). In particular, the main ventral hippocampal output region, the ventral subiculum (vS), is composed of multiple neuronal populations that send parallel, long-range projections to distinct areas, including prefrontal cortex (vS^PFC^), lateral hypothalamus (vS^LH^) and nucleus accumbens shell (vS^NAc^) (Naber and Witter, 1998). These populations are proposed to integrate a myriad of local and long-range inputs (Amaral and Cowan, 1980; Strange et al., 2014; Wyss et al., 1979) to perform their unique behavioural functions (Adhikari et al., 2010; Cembrowski et al., 2018a; Ciocchi et al., 2015; Jimenez et al., 2018; Soltesz and Losonczy, 2018). However, to date, knowledge of the input connectivity of these vS output neurons is lacking. Further, these vS populations are spatially patterned along the proximal-distal axis (ranging from the CA1 to the presubiculum borders) (Cembrowski et al., 2018a), and synaptic input varies dramatically across this axis (Aggleton and Christiansen, 2015; Cembrowski et al., 2018a; Knierim et al., 2014; Masurkar et al., 2017; van Groen et al., 2003). Based on this, we hypothesised that different vS output populations receive distinct upstream inputs, and reasoned that these inputs may in turn depend on the spatial location (Cembrowski et al., 2018a; 2016) or downstream target (Ciocchi et al., 2015; Jimenez et al., 2018; Naber and Witter, 1998) of postsynaptic neurons, or a combination of these two factors.

To address this hypothesis, we applied rabies tracing across different vS projections and proximal-distal postsynaptic cell locations. We obtained a brain-wide map of inputs to vS subpopulations and identified quantitative differences in multiple long-range input regions to vS that depended to different extents on the spatial location and projection target of vS neurons. One such input region was the nucleus reuniens (RE), a thalamic structure that is essential for hippocampal-dependent learning and memory functions, such as contextual learning (Xu and Sudhof, 2013) and goal-directed behaviour (Ito et al., 2015). The circuit model underlying these functions is proposed to be a reciprocal vS-PFC-RE-vS loop, where RE links the PFC and hippocampus (Dolleman-van der Weel et al., 2019; Vertes, 2006). Specifically, it is assumed that, within the hippocampus, vS^PFC^ neurons receive and integrate RE input and transmit these signals back to PFC. Unexpectedly, we found that RE inputs did not connect with vS^PFC^ neurons, and instead preferentially innervated vS^NAc^ and vS^LH^ neurons. We used *channelrhodopsin-(ChR2)- assisted circuit-mapping* (CRACM) to functionally validate the strength of RE inputs in vS and confirmed that - contrary to previous assumptions - RE input was functionally biased away from vS^PFC^ neurons.

## Results

### Hippocampal projection populations are distributed along the proximal-distal axis

The vS projection populations that target discrete downstream regions are non-overlapping and intermingled (Naber and Witter, 1998), and each population occupies a unique spatial distribution along the proximal-distal axis of the hippocampus (Kim and Spruston, 2011). As long-range input into hippocampus has been shown to vary dramatically across this axis (Aggleton and Christiansen, 2015; Cembrowski et al., 2018a; Knierim et al., 2014; Masurkar et al., 2017; van Groen et al., 2003), we first wanted to quantify the spatial distribution of the different projection populations within vS. We stereotaxically injected the retrograde tracer cholera toxin subunit-B (CTXβ) into PFC, LH or NAc in a pairwise manner (**Figure 1A**). We confirmed that vS^PFC^, vS^NAc^ and vS^LH^ neurons reside predominantly in the output region vS, and that the fraction of colocalised cells (neurons that projected to more than one injection site) ranged from 2 to 6% (Naber and Witter, 1998) (**Figure 1B-C**). By examining where each projection population was located along the proximal-distal axis, we observed that vS^PFC^ neurons tended to be positioned most proximally at the CA1/subiculum border, vS^LH^ neurons most distally in subiculum, while vS^NAc^ neurons were widely distributed and spanned the area bounded by both vS^PFC^ and vS^LH^ (**Figure 1D-F, Figure 1-figure supplement 1**). Notably, this spatial arrangement of vS^NAc^ differed from that observed in the dorsal subiculum (Cembrowski et al., 2018a), where a clear anatomical boundary separates projections to NAc and hypothalamus. Together, these data indicate that vS projection populations are segregated cell types and occupy overlapping yet distinct locations along the proximal-distal axis.

**Figure 1:**
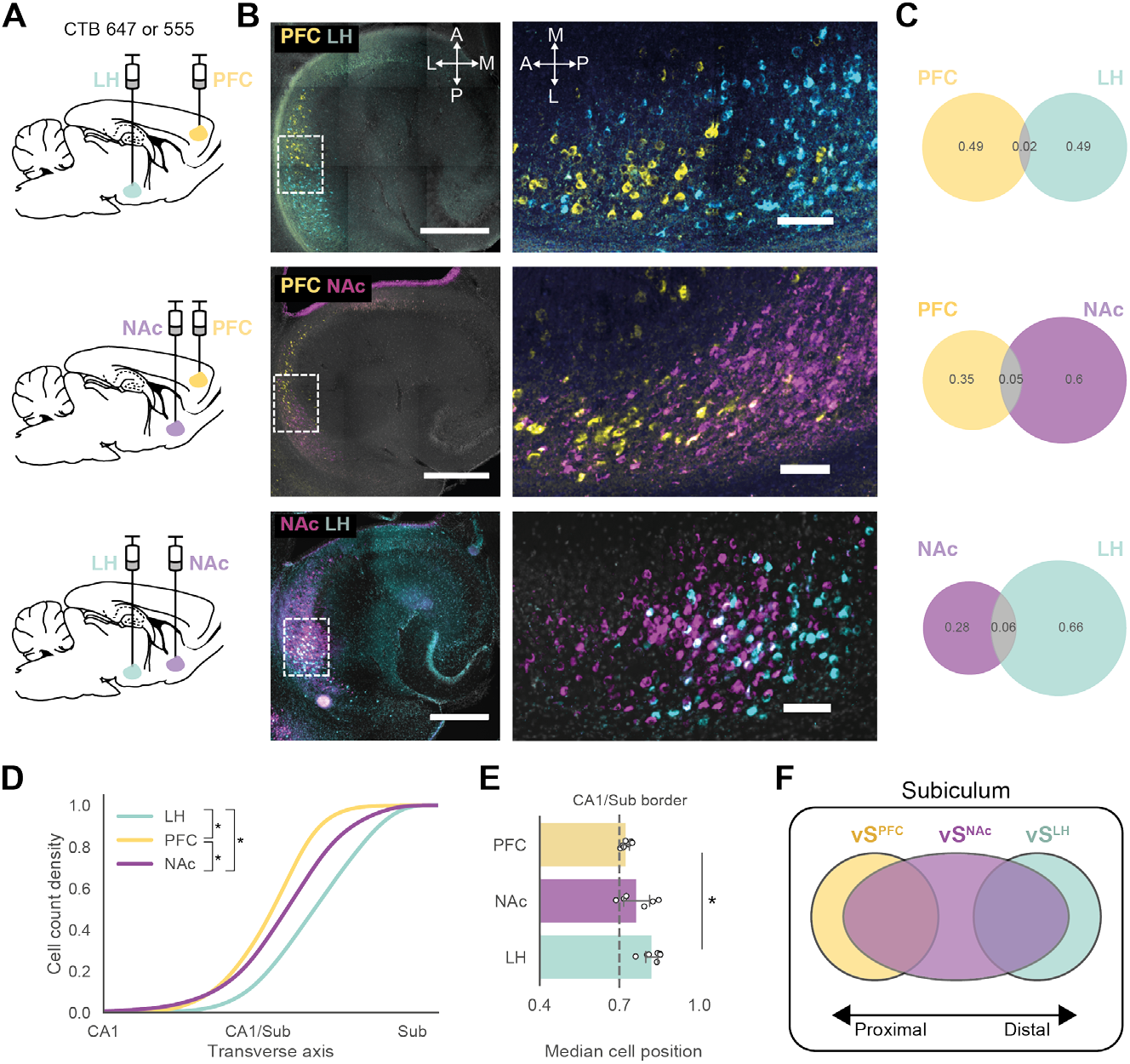
Non-overlapping vS neurons occupy distinct spatial sites. (A) Schematic of pairwise CTXβ injections into NAc, PFC or LH to retrogradely label vS neurons projecting to these areas. (B) *Left:* Example horizontal section images of the vS with retrogradely labelled cells. *Right:* Zoom-in images of the boxed area from *Left*, illustrating the vS region where most cells are labelled. Individual vS projections show distinct spatial patterns along the A-P axis. Scale bar: 500 μm (right) 100 μm (left). (C) Venn diagrams showing cell quantification from n = 6 – 8 sections per hemisphere from 3 – 4 hemispheres. Proportionally few cells (2 – 6%) are colocalised. (D) Kernel density plots of cell count density along the proximal-distal axis. The x-axis is a normalised range from 0 (dentate gyrus hilus) to 1 (end of subiculum), where CA1 corresponds to 0.4, CA1/Sub border to 0.7 and Subiculum to 1 along this normalised range. The data are plotted as cumulative distributions, and the spatial distributions of vS^PFC^, vS^NAc^ and vS^LH^ somata were different (pairwise two-sample Kolmogorov-Smirnov tests, adjusted for multiple comparisons with Bonferroni correction, see Extended Table 2). (E) Median cell positions of vS^PFC^, vS^NAc^ and vS^LH^ along the proximal-distal axis were distinct (n = 6 – 7 hemispheres per projection, one-way ANOVA, F_2,17_ = 9.56, p = 0.0017). (F) Schematic of findings in Figure 1, where vS^PFC^ neurons are located most proximally, vS^LH^ neurons most distally and vS^NAc^ neurons span the area between both vS^PFC^ and vS^LH^.

### Labelling of hippocampal input dependent on spatial location and projection target

Next, to directly assess the organisation of presynaptic inputs onto these vS projections, we applied *tracing the relationship between input-output* (TRIO) (Beier et al., 2019; 2015; Ren et al., 2018; Schwarz et al., 2015) to vS projections (**Figure 2A, Figure 2-figure supplement 1**). Briefly, this approach involved injecting *AAV2-retro-Cre* into the output target region to retrogradely express Cre recombinase in vS neurons that projected to the target site. In the same surgery, we injected a single Cre-dependent helper construct (*AA2/1-synP-FLEX-split-TVA-2A-B19G*, or TVA-G) into vS. After 2 weeks of TVA-G expression in projection-specific neurons, G-deleted, pseudotyped rabies virus (*EnvA-RV1JG-H2B-mCherry*) harbouring nuclear-localised mCherry was injected into vS to infect TVA-G+ cells. Importantly, we varied the injection site of TVA-G and rabies along the antero-posterior (A-P) axis - which approximates the proximal-distal axis in vS - to ensure that we sampled starter cells (i.e. cells from which rabies virus begins the monosynaptic retrograde tracing) from different proximal-distal vS locations (**Figure 2B, Figure 2-figure supplement 2)**. In parallel, we confirmed in control experiments that input labelling by rabies infection required the presence of both Cre and TVA-G (**Figure 2-figure supplement 3**). Our overall strategy thus provided us with experimental control over both starter cell location (by injection location of TVA-G and rabies within vS) and output projection (by injection location of *AA2V-retro-Cre* in target sites). This approach allowed us to assess how variations in spatial position and projection of vS neurons influence input size and identity.

**Figure 2:**
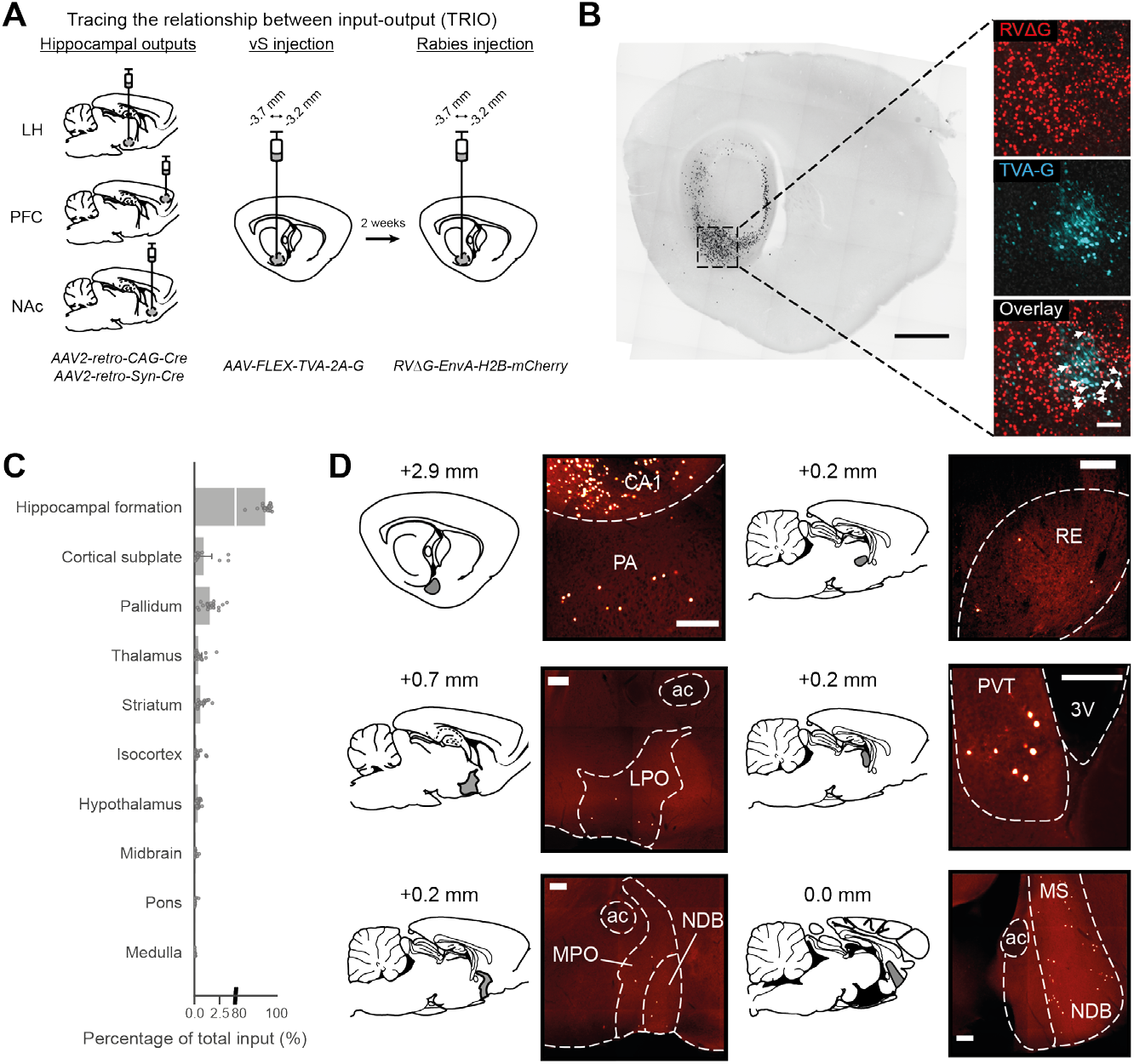
TRIO of vS output neurons. (A) Schematic of TRIO. (B) *Left:* Example sagittal section image of the brain with rabies-labelled cells (black) in the hippocampus. *Right:* Zoom-in images of the boxed area from *Left*, illustrating the vS region where starter cells are located. Starter cells are cells that colocalise for TVA-G (cyan) and rabies (red; arrows) and represent cells from which monosynaptic retrograde tracing begins. Scale bar: 1000 μm (right), 200 μm (left). (C) Bar plot illustrating coarse-level input-mapping normalised to the total number of inputs counted in a single brain. The majority of inputs arise from within the hippocampal formation with contributions from numerous extrahippocampal regions. Note the break in the x-axis. (n = 16 brain) (D) Representative subcortical long-range monosynaptic inputs to all vS neurons. Approximate sagittal planes are displayed with grey shaded boxes indicating the estimated locations of the corresponding images. The numbers above the sagittal plates are the distance (medial-lateral) from bregma. Scale bar: 200 μm. Error bars represent sem.

### Brain-wide rabies tracing of vS projection neurons

To quantify input to vS, we conducted brain-wide cell counts of rabies labelled neurons and registered the data to the Allen Brain Atlas (Fürth et al., 2018; Oh et al., 2014). Consistent with previous studies, the majority (~90%) of direct input to all vS projection types and spatial locations arose locally within the hippocampal formation (**Figure 2C**). In addition, there were numerous long-range inputs arising from ~38 brain regions spanning thalamic, striatal, pallidal, cortical, hypothalamic and amygdalar regions (**Figure 2C-D**) including the nucleus of diagonal band, medial septum, nucleus reuniens and posterior amygdala. We also consistently observed inputs arising from the preoptic area (both lateral and medial areas).

Next, we wanted to quantify the long-range input onto vS neurons depending on their projection target (PFC, LH or NAc), or spatial location along the proximal-distal axis (centre of mass, or COM, see Methods, **Figure 3A, Figure 3-figure supplement 1-3**). For each brain region that contributed more than 1% of extrahippocampal input (18 discrete regions), we built a linear regression model with both COM and projection information as predictors (**Figure 3B**). Using this strategy, we found that input from medial preoptic area (MPO), posterior amygdala (PA) and nucleus reuniens (RE) resulted in statistically significant model fits, indicating that these upstream regions provided quantitatively different input sizes to vS depending on either COM, output projection or both predictors.

**Figure 3:**
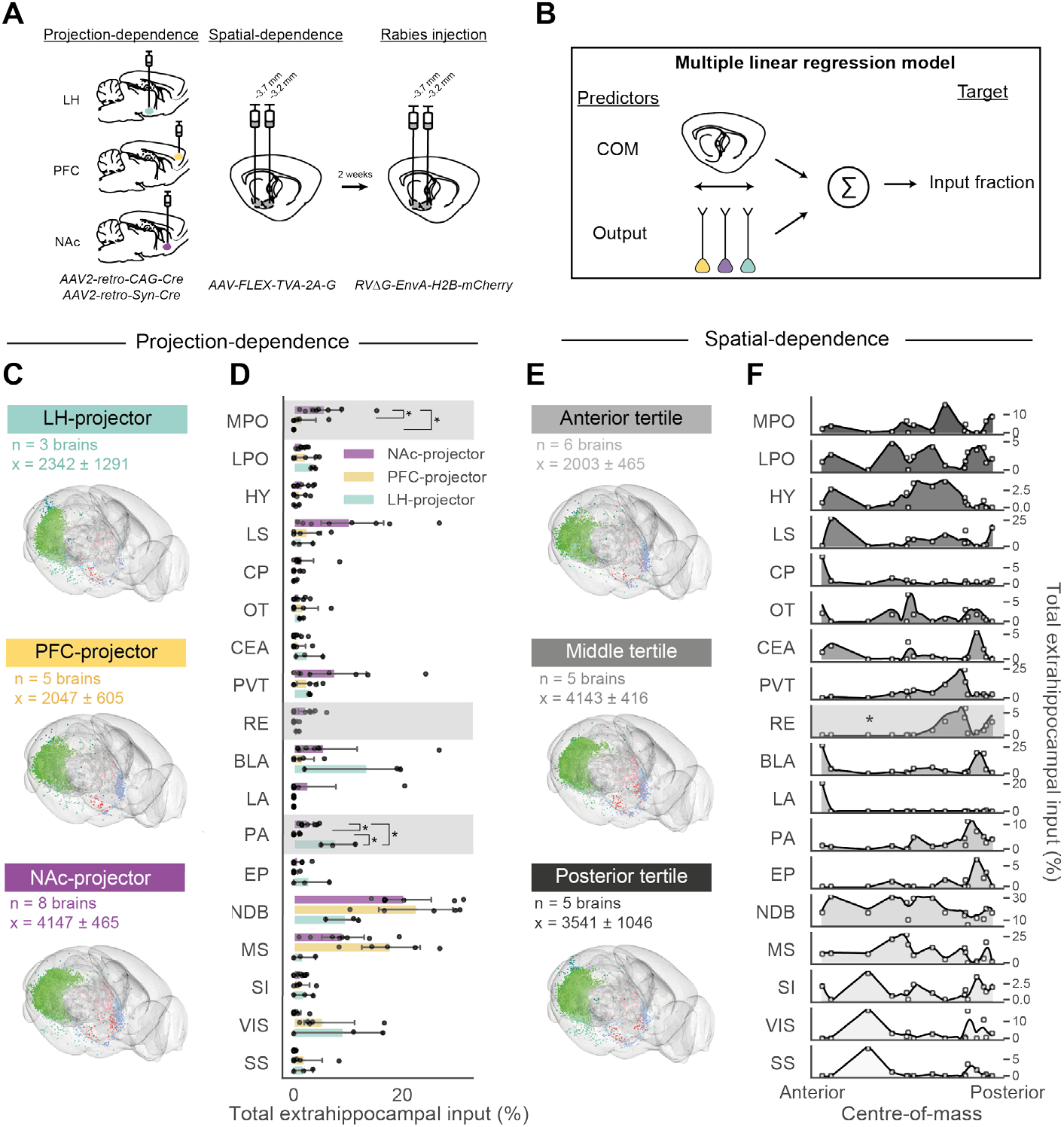
TRIO of vS output neurons reveals projection and spatial bias of inputs. (A) Schematic of TRIO protocol that allows the assessment of projection-dependence of inputs through retro-Cre injection in the output site, and spatial-dependence through different A-P injection sites of TVA-G and rabies. (B) Schematic of linear regression analysis. Linear models were constructed with a regressor matrix composed of COM and output type as regressors, and the dependent variable as the extrahippocampal input fraction. (C–F) Quantification of extrahippocampal long-range monosynaptic inputs. The number of inputs from a brain region is expressed as a percentage of the total number of extrahippocampal inputs (i.e. inputs from outside the hippocampal formation). The average cell numbers in (C) and (E) are expressed as mean ± sem. (C–D) Dataset split according to output projection. (C) Glass brain plots of inputs detected from all brain samples. (D) Bar plots of extrahippocampal input fractions split by projection populations. Shaded regions indicate input regions with significant multiple linear regression model fits as assessed by ANOVA and computing F-statistics (p < 0.05 after correction for multiple comparisons) and with significant projection-dependence (Wald’s test for model coefficients). Statistically significant model fits were followed up with post-hoc pairwise Tukey multiple comparisons (*p < 0.05). (E–F) Same dataset as in C and D but plotted as a function of standardised COM anterior-posterior (A–P) coordinates. For the glass brain plots in (E), the dataset and COM A-P coordinates were binned into tertiles along the A-P axis. (F) Input fractions as a function of COM. The shaded continuous line represents the smoothed input density (normalised with area under the curve = 1) as a function of COM. The input from RE (*) resulted in a significant model fit with a statistically significant COM coefficient.

We next investigated the relative contribution of projection type or COM by comparing single-predictor (projection *OR* COM) models with combined (projection *AND* COM) models (see Methods, **Figure 3-figure supplement 4A-B**). Using this approach, we found that MPO, PA and RE all innervated vS dependent on the projection target of the postsynaptic neuron (**Figure 3C-D**), while RE input was also dependent on starter cell location (**Figure 3E-F**). In summary: MPO input selectively targeted vS^NAc^ neurons; PA input selectively targeted vS^LH^ and vS^NAc^ neurons; while RE input targeted vS^LH^ and vS^NAc^ neurons only in distal locations within vS. As a control for our analysis, we tested the spatial- and projection-dependence of inputs separately and found that MPO and PA were projection-dependent while RE was spatially-dependent (**Extended Table 2**). Finally, we obtained a qualitative description of all intra- and extrahippocampal inputs (45 regions) and their relative dependence on COM or projection, which revealed a wide range in the relative dependence of different synaptic input on both COM and projection identity (**Figure 3-figure supplement 4C-D**). Overall, our rabies tracing dataset identified brain-wide regions that project to vS, including quantitatively biased inputs from MPO, RE and PA that depend differentially on the location and projection identity of the postsynaptic neuron.

### Biased nucleus reuniens input to hippocampal projections

A surprising finding in our dataset was that the RE input was anatomically biased to avoid vS^PFC^ neurons (**Figure 4A**). This finding runs counter to classic models of the PFC-RE-vS circuit where RE functions as a relay between PFC and hippocampus via long-range input to vS^PFC^ projections (Dolleman-van der Weel et al., 2019; Vertes, 2006). We thus sought to confirm this anatomical data using CRACM to ensure that this result was not due to methodological constraints such as viral tropism (Luo et al., 2018) or activity-dependence of viral tracing (Beier et al., 2017). From our tracing data, we hypothesised that the RE input was both spatially- and projection-biased, i.e. RE input targets distal areas where vS^PFC^ neurons are not abundant (COM), and also does not form synaptic connections with vS^PFC^ neurons in distal locations (Projection; see **Figure 3**). We injected AAV to express ChR2 in RE **(Figure 4B)**and found that ChR2+ axons emanating from RE were observed most densely in the distal region of vS (**Figure 4C**), consistent with the spatial dependence of our rabies tracing data. In these slices, we recorded light-evoked postsynaptic currents from pairs of retrogradely labelled neurons within axon-rich distal vS. By recording from pairs of neighbouring neurons within the same slice, this approach allowed us to directly compare the relative RE input strength across projection populations while controlling for proximal-distal position, ChR2 expression and light intensity. We observed that excitatory RE input was indeed much weaker onto vS^PFC^, whereas it strongly targeted neighbouring vS^LH^ and vS^NAc^ neurons (**Figure 4D-K, Figure 4-figure supplement 1**). Overall, these results demonstrate a functional bias of RE inputs away from vS^PFC^ and towards both vS^LH^ and vS^NAc^.

**Figure 4:**
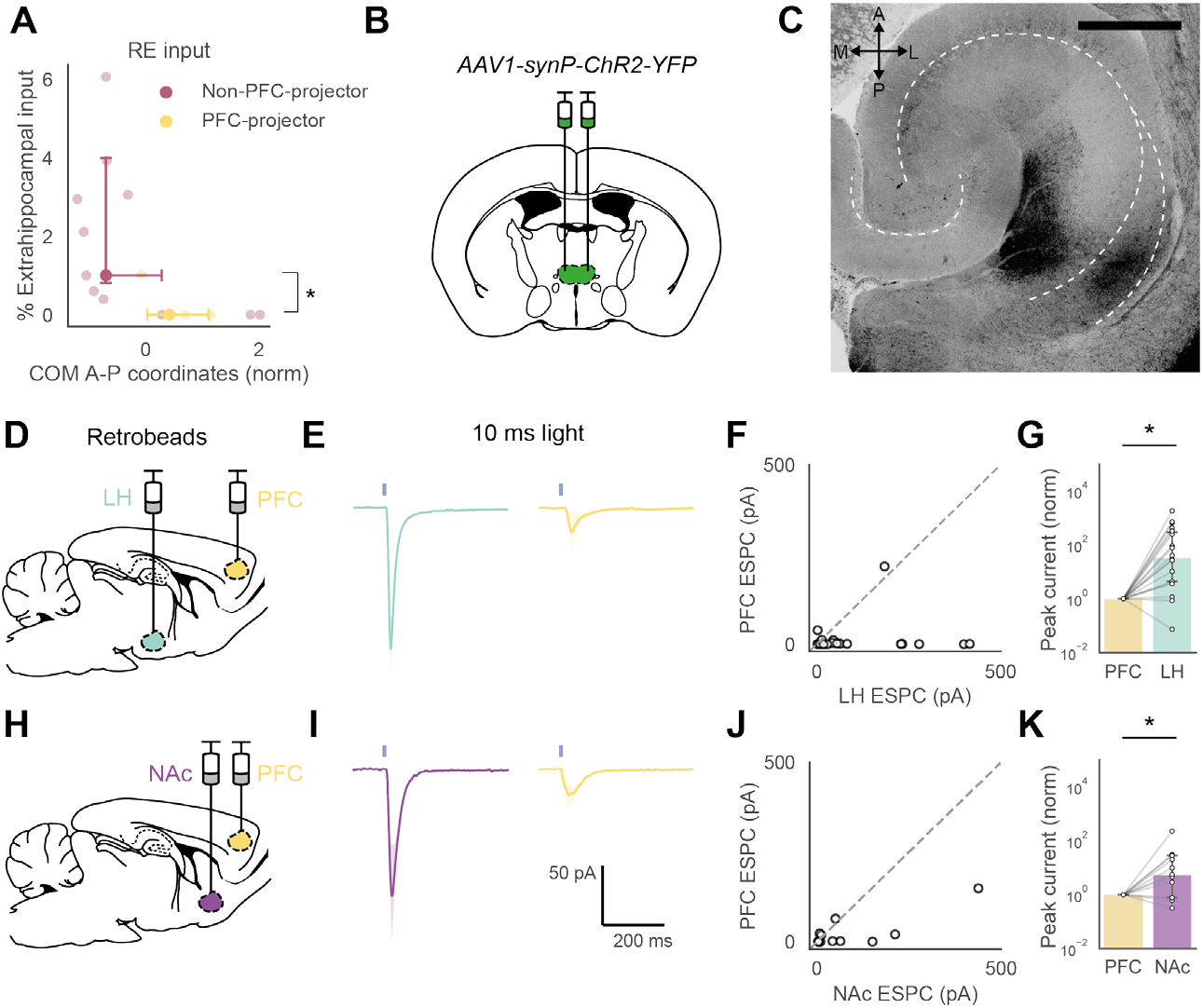
Nucleus reuniens inputs to vS projection neurons are functionally biased. (A) Rabies tracing monosynaptic inputs from RE as a function of COM and projection population. Non-PFC projectors (i.e. pooled vS^NAc^ and vS^LH^) have overall higher RE inputs than vS^PFC^ (n = 11 non-PFC projectors, n = 5 PFC-projectors, Mann-Whitney U test, U = 12.0, p = 0.038). Data expressed as percent input of extrahippocampal inputs counted in a single brain. Data points represent the median, error bars represent the inter-quartile range. (B) An AAV expressing ChR2 under the *synapsin* promoter was injected bilaterally into RE. (C) Confocal image of a horizontal section of hippocampus. RE axons are distributed near the CA1/subiculum border. (D, H) In the same surgery after ChR2 injection into RE, red and green retrobeads were injected into either NAc and PFC or LH and PFC to retrogradely label vS neurons that projected to the targeted site. (E, I) Light-evoked mean excitatory postsynaptic currents in pairs of (E) vS^LH^ and vS^PFC^, or (I) vS^NAc^ and vS^PFC^. All data represent the light-evoked response from 10 ms pulse of blue light. *Solid line*: mean photoresponse, *shaded region:* 95% confidence interval. (F, G) Scatter plots of light-evoked photocurrents from pairs. (G, K) Normalised EPSCs scaled to the photocurrent elicited in vS^PFC^, where the relative amplitudes of light-evoked photocurrents are higher in vS^LH^ (Wilcoxon signed-rank test, n = 18 pairs from 3 animals, V = 4.0, p = 3.86e-4) and in vS^NAc^ (Wilcoxon signed-rank test, n = 11 pairs from 3 animals, V = 6.0, p = 0.016). Bar plots in G and K indicate median and error bars indicate 95% bootstrapped confidence interval. *p < 0.05.

**Figure 5:**
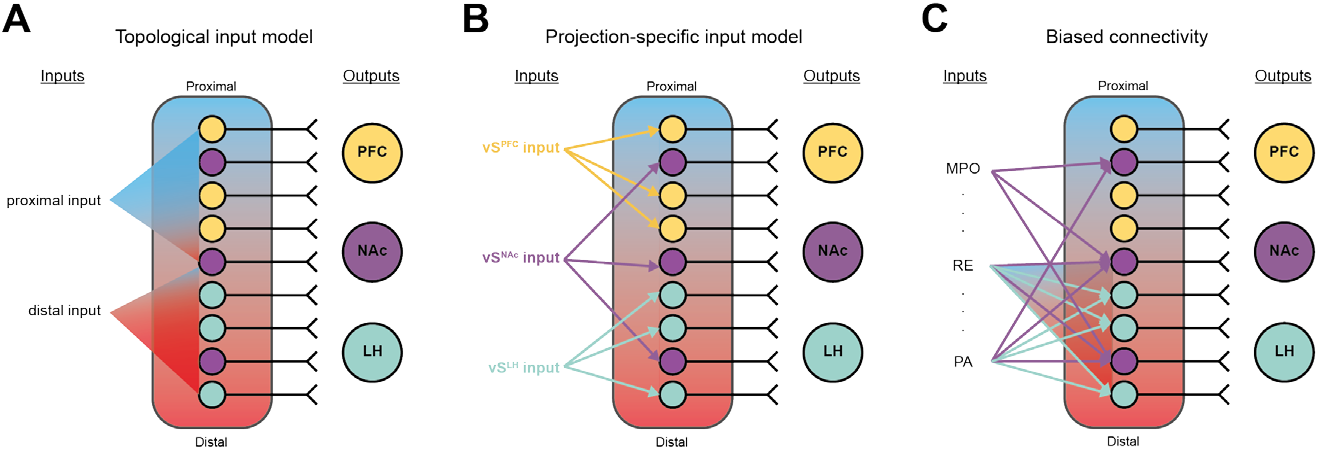
Topological and projection-defined input connectivity of vS output neurons. (A) Models of vS input connectivity based on the spatial location or (B) projection-specificity of postsynaptic vS neurons. (C) Our dataset supports a model of vS input-output connectivity in which there are co-existing inputs that are projection-, COM- and both projection- and COM-biased. In Figure 3, we found that MPO and PA inputs are projection-biased, where MPO input preferentially targets vS^NAc^ and PA input tends to connect with vS^NAc^ and vS^LH^. RE is both projection- and COM-biased, where inputs are spatially biased towards distal vS regions and target both vS^NAc^ and vS^LH^.

## Discussion

In this study, we leveraged the parallel nature of vS projections (Naber and Witter, 1998) and their unique spatial pattern to investigate the input organisation onto projection-defined vS populations. Using TRIO, we demonstrated that the organisation of inputs to vS output circuitry depended to different degrees on the spatial position and projection target of the postsynaptic neuron. Of the identified long-range inputs, MPO, RE and PA gave rise to quantitatively different inputs, where MPO selectively innervated vS^NAc^, PA selectively innervated vS^NAc^ and vS^LH^, and RE innervated vS^NAc^ and vS^LH^ only in distal vS. To our knowledge, the consistent MPO inputs to vS that we found have not been previously described in the literature (Wyss et al., 1979). These identified inputs also complement recent theories about the role of vS and its upstream circuitries. For example, the biased MPO input to vS^NAc^ may be important in mediating social reward (McHenry et al., 2017; Okuyama et al., 2016).

A surprising result from our dataset was that the RE input did not innervate vS^PFC^ neurons (**Figure 3, Figure 4**). The RE is essential for bidirectional communication between hippocampus and PFC. This thalamic region is proposed to form an anatomical link between hippocampus and PFC, thereby closing a PFC-RE-vS-PFC loop (Dolleman-van der Weel et al., 2019; Ito et al., 2015; Vertes, 2006). Little information is present about the extent to which the projection-defined vS populations are involved in this circuit loop, and it is generally assumed that RE input is integrated by vS^PFC^ neurons to relay signals from hippocampus to PFC. By demonstrating that vS^PFC^ receive anatomically few and functionally weaker RE inputs, our data challenge this circuit model, and instead suggest that RE input is integrated upstream of vS^PFC^ neurons - i.e. either through dorsal hippocampus, entorhinal cortex, local interneurons or local pyramidal cell populations - before being transmitted back to PFC. Thus, it will be necessary for future work to investigate in more detail the hippocampal microcircuitry that is involved in integrating RE input, as well as the details of other multi-synaptic routes that may allow reciprocal connectivity between the hippocampus and PFC, such as via entorhinal cortex and amygdala.

Interestingly, we also show that RE input is anatomically and functionally biased towards vS^NAc^ and vS^LH^ neurons. This finding suggests that RE does not only communicate with hippocampus and PFC but is embedded within a wider network that encompasses limbic regions such as striatum and hypothalamus (Dolleman-van der Weel et al., 2019). This observed circuit connectivity further complements previous data that demonstrated a crucial role for RE in goal-directed planning (Ito et al., 2015) and fear generalisation (Xu and Sudhof, 2013) - functions that may be crucial for the proposed roles of vS^NAc^ in reward-seeking and vS^LH^ in anxiety (Cembrowski et al., 2018a; Ciocchi et al., 2015; Jimenez et al., 2018). Therefore, it will be important to further elucidate the role of RE input in these vS-related behaviours. This is particularly important given the key role of this circuit in preclinical models of disorders such as schizophrenia, depression and Alzheimer’s disease (Dolleman-van der Weel et al., 2019; Ito et al., 2015; Vertes, 2006).

Finally, our overall dataset supports a model of combined topological and output-defined connectivity of vS inputs (**Figure SA-C**), where depending on the upstream region, rabies-labelled inputs are biased according to space, projection type or both these factors. This is in keeping with the known spatial and projection-specific specialisations of subiculum; across proximal-distal subdivisions, proximal vS is involved in sensory encoding (Knierim et al., 2014) and distal vS supports path integration (Cembrowski et al., 2018a; Knierim et al., 2014), while across projection populations, vS^PFC^ and vS^LH^ encode innate threat (Adhikari et al., 2010; Ciocchi et al., 2015; Jimenez et al., 2018) and vS^NAc^ encodes social memory (Okuyama et al., 2016). Crucially, the joint spatial- and projection-dependence of RE inputs also support the existence of spatial- and projection-specific function. For example, goal-directed locomotion has been proposed to be both specific to vS^NAc^ neurons and distal subiculum (Cembrowski et al., 2018a; Ciocchi et al., 2015; Okuyama et al., 2016). Investigating how the identified inputs are functionally coupled to each of these vS subpopulations, and how different behaviours recruit these upstream circuitries, will be a goal for future experiments. Overall, our study has revealed a basis for the selective control of individual vS projection neurons, through a biased organisation in the brain-wide input connectivity of the vS parallel circuitry.

## Materials and Methods

### Animals

Young adult male mice (CTB and rabies tracing: at least 7 weeks old; physiology: 7 - 9 weeks old) provided by Charles River were used for all experiments. All animals were housed in cages of 2 to 4 in a temperature- and humidity-controlled environment with a 12 h light-dark cycle (lights on at 7 am to 7 pm). All experiments were approved by the UK Home Office as defined by the Animals (Scientific Procedures) Act, and University College London ethical guidelines.

### Retrograde tracers

Red and green fluorescent retrobeads (Lumafluor Inc.) were used for electrophysiological recordings. Cholera toxin subunit B (CTXβ) tagged with Alexa 555 or 647 (Molecular Probes) were used for retrograde tracing.

### Viruses

|  |  |
| --- | --- |
| AAV2-retro-CAG-Cre | (UNC Vector Core) |
| pENN-AAV-hSyn-Cre-WPRE-hGH | (gift from James M. Wilson, Addgene viral prep # 105553-AAV1) |
| AA2/1-synP-FLEX-split-TVA-EGFP-B19G | (a gift from Ian Wickersham, Addgene viral prep #52473, see ref) |
| EnvA-RVΔG-H2B-mCherry-2A-CLIP | (a gift from Marco Tripodi, MRC LMB, unpublished) |
| AAV1-synP-ChR2-YFP | (a gift from Karl Deisseroth, Addgene viral prep # 26973-AAV1) |

### Stereotaxic surgery

Stereotaxic surgeries were performed according to previously described protocols (Cetin et al., 2007). Mice were anaesthetised with isoflurane (4% induction, 1.5 to 2% maintenance) and secured onto a stereotaxic apparatus (Kopf). A single incision was made along the midline to reveal the skull. AP, ML and DV were measured relative to bregma, and craniotomies were drilled over the injection sites. Long-shaft borosilicate pipettes were pulled and backfilled with mineral oil, and viruses were loaded into the pipettes. Viruses were injected with a Nanoject II (Drummond Scientific) at a rate of 13.8 nL every 10s. Following infusion of the virus, the pipette was left in place for an additional 10 mins before being slowly retracted. The following coordinates (in mm) were used:

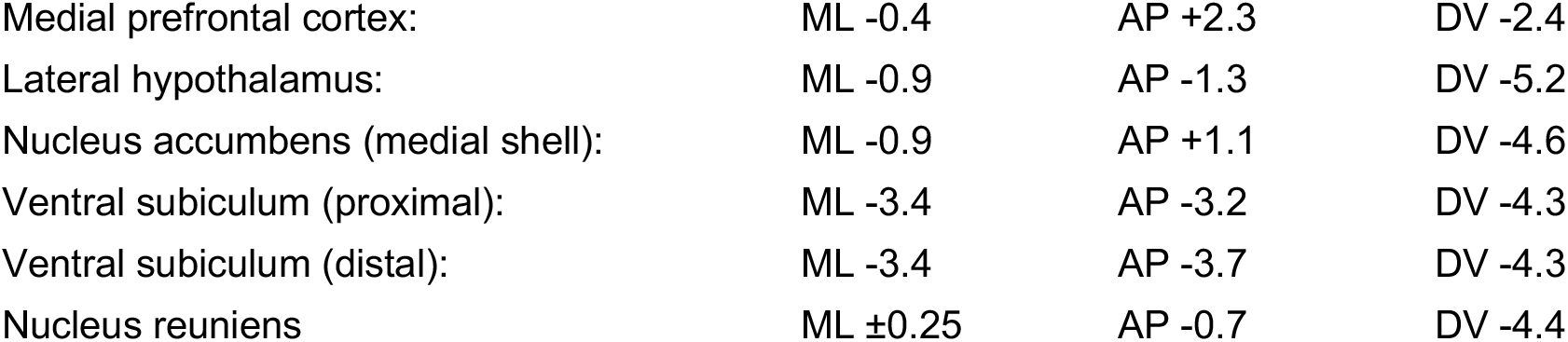

|  |  |  |  |
| --- | --- | --- | --- |
| Medial prefrontal cortex: | ML -0.4 | AP +2.3 | DV -2.4 |
| Lateral hypothalamus: | ML -0.9 | AP -1.3 | DV -5.2 |
| Nucleus accumbens (medial shell): | ML -0.9 | AP +1.1 | DV -4.6 |
| Ventral subiculum (proximal): | ML -3.4 | AP -3.2 | DV -4.3 |
| Ventral subiculum (distal): | ML -3.4 | AP -3.7 | DV -4.3 |
| Nucleus reuniens | ML ±0.25 | AP -0.7 | DV -4.4 |

Following injection of substances into the brain, animals were sutured and recovered for 30 mins on a heat pad. Animals received carprofen as a peri-operative s.c. injection (0.5 mg / kg) and in their drinking water (0.05 mg / mL) for 48 hours post-surgery.

### Retrograde tracing

For CTXβ retrograde tracing, 150 nL of Alexa 555- or 647-tagged CTXβ was injected into one of three output regions (PFC, NAc or LH). After at least 14 days post-surgery, animals were sacrificed for histology. For rabies monosynaptic tracing, experiments were done according to previously described protocols (Beier et al., 2015). Adult male mice were injected with 100 nL of *AAV2retro-CAG-Cre* or *AAV2retro-synP-Cre* into one of three output regions (PFC, NAc or LH), and in the same surgery, 250 nL of *AA2/1-synP-FLEX-split-TVA-EGFP-B19G* was injected into ventral subiculum (proximal) or ventral subiculum (distal). In a second surgery 2 weeks later, 300 - 400 nL of *EnvA-RV1JG-H2B-mCherry* was injected into vS. After 7 days of rabies expression, animals were sacrificed for histology.

### Histology and imaging

Animals were deeply anaesthetised with a lethal dose of ketamine and xylazine (100 mg/kg) and perfused transcardially with phosphate-buffered saline (pH = 7.2) followed by 4% paraformaldehyde. Brains were dissected and post-fixed overnight at 4°C prior to sectioning. For CTXβ tracing analysis, 60-μm sections in the horizontal plane were prepared using a vibratome (Campden Instruments). For brain-wide rabies tracing analysis, 60-μm sections in the sagittal plane were prepared in sagittal sections with a supporting block of agar, and every 2^nd^ section was mounted. Sections were mounted on Superfrost Plus slides with ProLong Glass antifade mounting medium (Molecular Probes) and imaged with a 5x objective using a Zeiss Axioscan Z1, using standard filter sets for excitation/emission. Images were exported as 16-bit depth files and analysed using ImageJ and WholeBrain software (Fürth et al., 2018).

### Analysis of spatial positions of vS projection populations

The spatial positions of retrogradely labelled vS projections to PFC, NAc and LH were analysed using ImageJ. 6 to 8 sections spanning vS (DV −3.5 to −4.5 mm) were analysed per hemisphere. Horizontal sections of the hippocampus were digitally straightened from the dentate gyrus to the end of subiculum using the *Straighten* function on ImageJ to approximate the proximal-distal axis. Labelled cells in each slice were manually counted and registered to this axis using the ImageJ CellCounter plugin. Each registered cell was collapsed in the radial axis (y-coordinate), and only the proximal-distal axis (x-coordinate) of each cell was used for spatial position analysis. The CA1/subiculum border occurs approximately at 0.7 within this normalised proximal-distal axis range and was anatomically defined as the disappearance of stratum oriens and the fanning out of the pyramidal cell layer. The spatial positions were analysed with custom Python routines.

### Brain-wide mapping and analysis of rabies-labelled inputs

Cell counting of rabies labelled inputs was conducted using Wholebrain (Fürth et al., 2018), a recently developed automatic segmentation and registration workflow in R. After acquiring the imaged sections and exporting as 16-bit depth image files, images in the rabies mCherry channel were manually assigned a bregma coordinate (ML −4.0 to 0.0 mm) and processed using WholeBrain (Fürth et al., 2018) and custom cell counting routines written in R. The workflow comprised of (1) segmentation of cells and brain section, (2) registration of the cells to the Allen Brain Atlas and (3) analysis of anatomically registered cells. As tissue section damage impairs the automatic registration implemented on the WholeBrain platform, sections with poor registration were manually registered to the atlas plate using corresponding points to clear anatomical landmarks. Once all cells have been registered, the cell counts were further manually filtered from the dataset to remove false-positive cells (e.g. debris). Virtually all cells were detected in the injected hemisphere, apart from a consistent set of contralateral CA3 inputs. Therefore, we only used the injected hemisphere up to the midline for cell quantification analysis.

Each cell registered to a brain region was classified according to the Allen Brain Atlas (ABA) brain structure ontology. Information on the ABA hierarchical ontology was scraped from the ABA API (link: http://api.brain-map.org/api/v2/structure_graph_download/1.json) using custom Python routines. Subsequently, we defined *a priori* the coarse-level (or parent) structures to which each cell belonged, which comprised the following: *Hypothalamus, lsocortex, Hippocampal formation, Thalamus, Cortical subplate, Pallidum, Striatum, Midbrain, Pons* and *Medulla.* Fine-level (or child) structures represent all brain regions existing as subcategories of the corresponding coarse-level structure, e.g. *nucleus reuniens* and *paraventricular thalamus* are fine-level (child) structures relative to *Thalamus* (parent). For quantification of input fractions, cells residing in different layers within the same structure, e.g. CA1 stratum oriens and stratum lacunosum-moleculare, or subdivisions of nuclei, e.g. basomedial amygdala, posterior division (BMAp) and anterior division (BMAa), were collapsed across layers and subdivisions and counted as residing in one single region (BMA). Note that lateral and medial entorhinal cortex (LEC and MEC, respectively) were exceptions to this rule, and we analysed MEC and LEC as separate structures.

### Starter cell centre of mass (COM) quantification

To determine the starter cell COM, every 2^nd^ sagittal section from both the rabies mCherry+ and TVA-G GFP+ channels that spanned the extent of the TVA-G GFP+ expression was obtained and analysed. Images were collected in order from lateral to medial. Colocalised cells representing starter cells were manually registered onto digital plates from the Paxinos atlas using the ImageJ ROI Manager function. Cells were collapsed in the ML and DV planes as the main source of spatial variation occurred in the AP plane, and the mean x-coordinates (corresponding to the AP dimension) represented the geometric COM. As the TVA-G construct expressed TVA and G bicistronically (i.e. the exon for TVA and G are linked by a self-cleaving 2A peptide), all cells were assumed to express TVA and G in a 1:1 stoichiometry and unlikely to express one gene without the other. Therefore, all colocalised cells were treated as starter cells.

### Analysis of COM vs. projection dependence of rabies-labelled inputs

For Figure 2, the input fraction was normalised to the total extrahippocampal inputs in the same brain. The dataset containing cell counts from n = 16 brains were analysed according to projection or COM. Only fine-level structures exceeding 1% of extrahippocampal input were assessed (18 brain regions). For projection-dependence analysis, 3D glass brains were plotted using the *glassbrain* function from the WholeBrain package split by vS projection. For spatial-dependence analysis, glass brains were plotted as tertiles of the COM in the A-P axis. The input density for each brain region as a function of COM was visualised by first sorting the input fractions of each brain by the COM in the posterior to anterior direction. The array of input fractions was then interpolated to produce 500 data points, smoothed with a Savitzky-Golay filter (window size = 51 points, order = 3), and normalised by dividing each data point by the total area under the curve. All *p* values generated from multiple comparisons were corrected using the Benjamini-Hochberg method (false-discovery rate < 0.05).

Multiple linear regression modelling was performed to compare the relative influence of COM and projection identity on the amount of rabies labelled inputs in a brain region of interest. For each brain region, a multiple regression model - in which the predictor variables were the COM and projection identity - was constructed using the *ols* function from the *statsmodel* package in Python. The overall statistical significance of full models (containing COM and projection as predictors) were assessed by ANOVA and computing the F-statistic values per brain region. The models with p-value < 0.05 were further analysed for statistically significant coefficients using the Wald test and followed up with post-hoc pairwise Tukey multiple comparisons between vS projection populations. To further assess the importance of each predictor to the model, the likelihood ratio (LR) test was used to compare the full model to a reduced model containing only one predictor - either COM or projection. The reduced models containing either COM or projection identity as a predictor were built using the same *ols* function. The LR test was computed using the following formula from the *statsmodel* package function *compare_lr_test:*

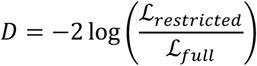

where 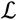 is the likelihood of the model, and D is a test statistic that follows a *χ*^2^ distribution with degrees of freedom (df) equal to the difference in the number of predictors between the full and reduced model. The p-values generated from multiple LR tests were corrected using the Benjamini-Hochberg method (FDR < 0.05). As a complementary analysis, the projection-dependence of extrahippocampal inputs from Fig 2F was further analysed using multiple one-way ANOVAs, while the spatial-dependence of these inputs was assessed using multiple Spearman rank correlation tests of input fractions against starter cell COM. This analysis revealed similar patterns of biased connectivity to the multiple linear regression analysis (**Extended Table 2**), where MPO and PA were detected as projection-dependent inputs, but RE was detected as only COM-dependent. This statistical result for RE was likely due to the fact that the multiple one-way ANOVA testing failed to take into account the combined effects of COM- and projection-dependence of RE inputs.

To assess the relative goodness-of-fit of non-nested *COM models* or *projection models*, we performed linear regression models by using the *ols* function from the Python package *statsmodels*. Linear models were built for each brain region, where the target variable was the input fraction observed in that brain region normalised to the total number of inputs, and the predictor variable was either the COM or projection. For each of 45 brain regions in total, there were 16 observations (n = 16 brains). The models were fitted, and goodness-of-fit was measured using the Bayesian information criterion (BIC) and adjusted R^2^. The BIC was computed using the following formula:

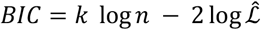

where *k* is the number of model parameters, *n* is the observation number and 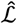 is the maximum likelihood of the model. The adjusted R^2^ was computed using the following formula:

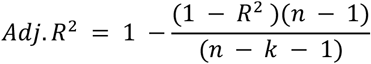

where *R*^2^ is the unadjusted R^2^ value computed as 1 - SSE/SST, *n* is the observation number and *k* is the number of model parameters. Finally, to compare model fits, the corresponding BIC and adjusted R^2^ values computed from COM and projection models were subtracted to obtain ΔBIC and ΔR^2^, respectively.

### CRACM experiments

#### Surgeries

CRACM of projection-defined vS neurons was done according to previously described protocols (MacAskill et al., 2014). 7- to 9-week old animals were injected with 250 nL of red (1:10 dilution) or green (undiluted) retrobeads in a counterbalanced order into two of three output regions (PFC, NAc or LH). In the same surgery, 250 nL of *AAV1-synP-ChR2-YFP* was injected into RE bilaterally. After at least 14 days of ChR2 expression, animals were sacrificed for electrophysiological recording.

#### Slice preparation

Acute, transverse hippocampal slices were used for all electrophysiological recordings. Mice were deeply anaesthetised with a lethal dose of ketamine and xylazine (100 mg / kg), and perfused transcardially with ice-cold sucrose solution containing (in mM): 190 sucrose, 25 glucose, 25 NaHCO_3_, 1.2 NaH_2_PO_4_, 10 NaCl, 2.5 KCl, 1 Na^+^ ascorbate, 2 Na^+^ pyruvate, 7 MgCl_2_ and 0.5 CaCl_2,_ bubbled continuously with 95% O_2_ / 5% CO_2_. Following perfusion, mice were decapitated, and their brains were rapidly dissected. The dissected brains were then placed in ice-cold sucrose solution and hemisected. The cerebellum was removed, and transverse slices were prepared using a vibratome (VT1200S, Leica), with a ~10° angle along the ventromedial plane to obtain sections that were orthogonal to the long-axis of the hippocampus. The thickness of hippocampal sections was 300 μm. Slices were transferred to a bath containing artificial cerebrospinal fluid (aCSF) and recovered first for 30 mins at 37°C, and subsequently for 30 mins at room temperature. The aCSF solution contained (in mM): 125 NaCl, 2.5 KCl, 1.25 NaH_2_PO_4_, 22.5 glucose, 1 Na^+^ ascorbate, 3 Na^+^ pyruvate, 1 MgCl_2_, 2 CaCl_2_. All recordings were performed at room temperature (22 – 24°C). All chemicals were from Sigma or Tocris.

#### Whole-cell electrophysiology and optogenetics

Whole-cell recordings were performed on retrogradely labelled hippocampal pyramidal neurons with retrobeads and visualised by their fluorescent cell bodies and targeted with Dodt contrast microscopy. For sequential paired recordings, neighbouring neurons were identified within one or two fields-of-view using a x40 objective at the same depth into the slice. The recording order of neuron pairs was counterbalanced to avoid complications due to rundown. Borosilicate recording pipettes (3 – 5 MO) were filled with a Cs-gluconate internal solution containing (in mM): 135 Gluconic acid, 10 HEPES, 10 EGTA, 10 Na-phosphocreatine, 4 MgATP, 0.4 Na_2_GTP, 10 TEA and 2 QX-314. Presynaptic glutamate release was elicited by illuminating ChR2 expressed in the presynaptic terminals of long-range inputs into the slice, as previously described (MacAskill et al., 2014). Wide-field illumination was achieved through a 40x objective with brief (0.2, 0.5, 1, 2, 5 and 10 ms) pulses of blue light from an LED centred at 473 nm (CoolLED pE-4000, with corresponding excitation-emission filters). Light intensity was measured as 4 - 7 mW at the back aperture of the objective and was constant between all recorded cell pairs. In all experiments, the aCSF contained 1 μM TTX and 100 μM 4-AP to isolate monosynaptic connectivity and increase presynaptic depolarisation, respectively. Recordings were conducted using a Multiclamp 700B amplifier (Axon Instruments), and signals were low-pass filtered using a Bessel filter at 1 kHz and sampled at 10 kHz. Data were acquired using National Instruments boards and WinWCP (University of Strathclyde) and analysed using custom routines written in Python 3.6.

For synaptic connectivity analysis, six recording sweeps were obtained for each optical pulse duration. The signals were preprocessed by baselining the signals to the first 100 ms, low-pass filtering using a Bessel filter (cutoff = 2 Hz, order = 2), averaging across the six recording sweeps for each optical pulse duration, and decimating the averaged signal to 1 kHz. The peak amplitude response of light-evoked EPSCs was measured as averages over a 2-ms time window around the peak compared to a 2-ms baseline period preceding the optical pulse. Only paired data in which at least one cell received > 5 pA were included for analysis for Fig 4D-K, while all cells irrespective of connectivity were included for the analysis described in Supplementary Fig 9. For each recorded neuron, the spatial position was obtained by manually registering the cell to a digital atlas depicting a horizontal section of the vS (−4.0 DV relative to bregma). The long axis of the CA1 and subiculum area (i.e. the y-coordinates) of each registered cell was used to approximate the anterior-posterior position. Input connectivity was determined by a threshold light-evoked response of > 5 pA.

#### Statistical analysis

All statistics were calculated using Python’s *scipy* and *statsmodels* packages, and R. Summary data are reported throughout the text and figures as mean ± sem unless otherwise stated. For Figure 1, normality of data distributions was determined by visual inspection of the data points. For Figure 2, the rabies tracing dataset was assessed for normality with Jarque-Bara test and equal variance with Levene’s test. The input fractions for multiple regions showed non-normality and heteroscedasticity in the input fractions across projection populations. Therefore, all analysis for Figure 2 were conducted after log-transformation of input fractions. Distributions of the data became more Gaussian-like after log-transformation as assessed again by the Jarque-Bara test and Levene’s test. All data were analysed using statistical tests described in the figure legends and in Extended Table 2. The alpha level was defined as 0.05. No power analysis was run to determine sample size *a priori*. The sample sizes chosen are similar to those used in previous publications.

## Acknowledgements

We thank Marco Tripodi and Fabio Morgese for the unpublished pseudotyped rabies virus, and Francesca Cacucci for assistance with surgery. We are grateful to Larry Swanson, Andy Murray, Tara Keck, Kenneth Harris and the MacAskill lab for helpful comments on the manuscript.

## Additional Information

### Competing lnterests

The authors declare that no competing interests exist.

### Author contributions

RWSW, Conception and design, Data acquisition, Analysis and Data Interpretation, Manuscript preparation. AFM, Conception and design, Data acquisition, Analysis and Data Interpretation, Manuscript preparation.

### Ethics

Animal experimentation. This study was approved by the UK Home Office as defined by the Animals (Scientific Procedures) Act and conducted in strict accordance with University College London ethical guidelines. All surgery was performed under isoflurane anaesthesia, and all steps were taken to minimise pain during procedures.

### Availability of Data and Software

Program code, scripts for statistical tests and original data are available on Github.

### Funding

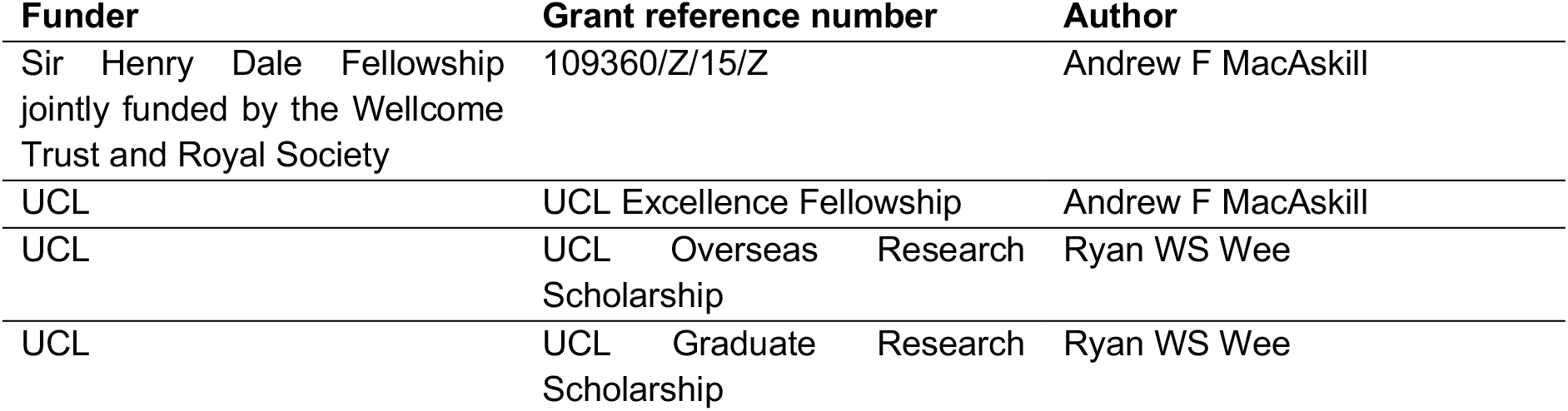

## Supplementary Figures

**Figure 1-figure supplement 1:**
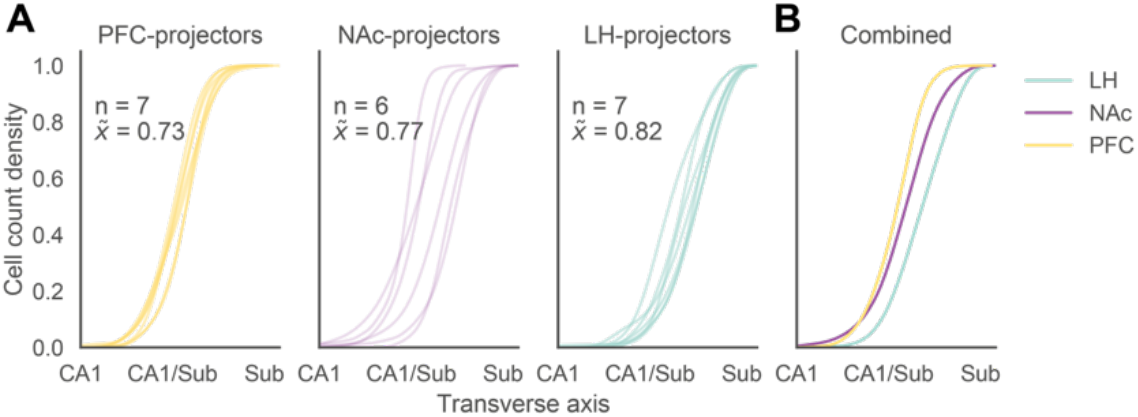
Distribution of CTB labelled cells by projection population. (A) CTXβ-labelled cells split by projection. Individual lines represent data of cell count distributions from one hemisphere. The number of hemisphere samples (n) and the average median position 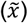 of the cell count distributions are provided within each subpanel. The x-axis is the normalised range of the transverse (proximal-distal) axis, and each hippocampal field was operationally defined as the following: CA1 is 0.4, CA1/Sub is 0.7 and Sub is 1 along the normalised transverse axis range. (B) Overlain plots of the pooled cumulative distribution for each vS projection. While mostly intermingled, each vS projection population tends to occupy distinct spatial sites.

**Figure 2-figure supplement 1:**
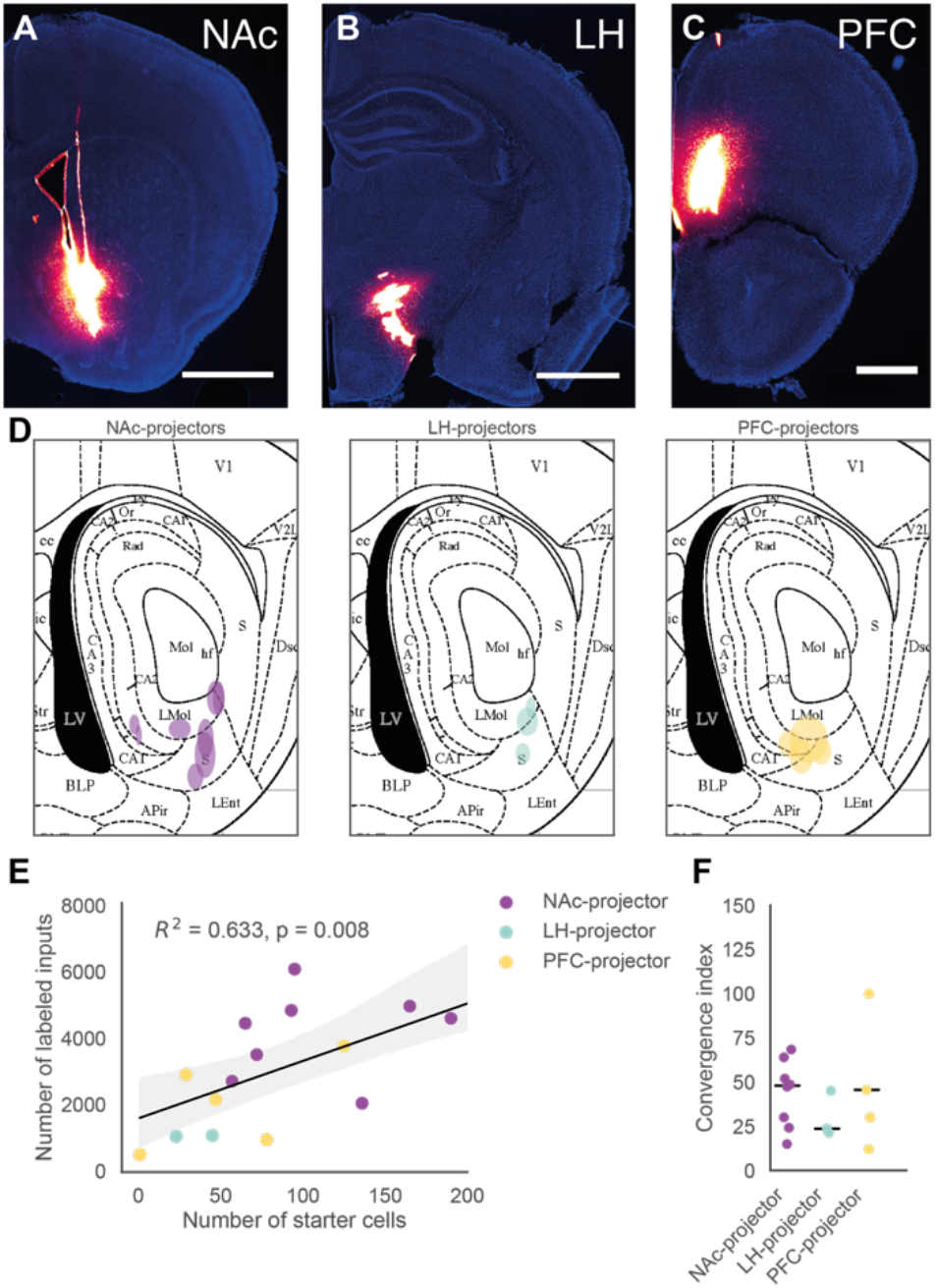
Starter cell quantification. (A – C) Example injection sites using identical volumes of CTXβ-647 and stereotactic coordinates as for *AAV2-retro-Cre* injections. All injection sites were localised in the targeted regions. Scale bar: 1000 μm (A-B), 500 μm (C). (D) Starter cell centre-of-mass (COM) plotted onto a reference plate from the Paxinos atlas; each ellipse represents the starter cell geometric mean from one brain sample (horizontal and vertical widths represent 1 s.d. about the mean coordinate position). vS^LH^ neurons are located more posteriorly (nearer to subiculum), vS^PFC^ neurons are positioned more anteriorly (nearer to CA1) while vS^NAc^ neurons span the regions occupied by both vS^LH^ and vS^PFC^ neurons. (E) Scatter plot of the number of starter cells against the total number of labelled inputs counted in each brain sample. The number of labelled inputs scales with the number of starter cells (Pearson correlation, R^2^ = 0.572, p = 0.021). Shaded regions represent 95% confidence intervals. (F) Convergent indices (the total number of inputs divided by the number of starter cells) plotted for each projection population. The solid line indicates the median convergent index.

**Figure 2-figure supplement 2:**
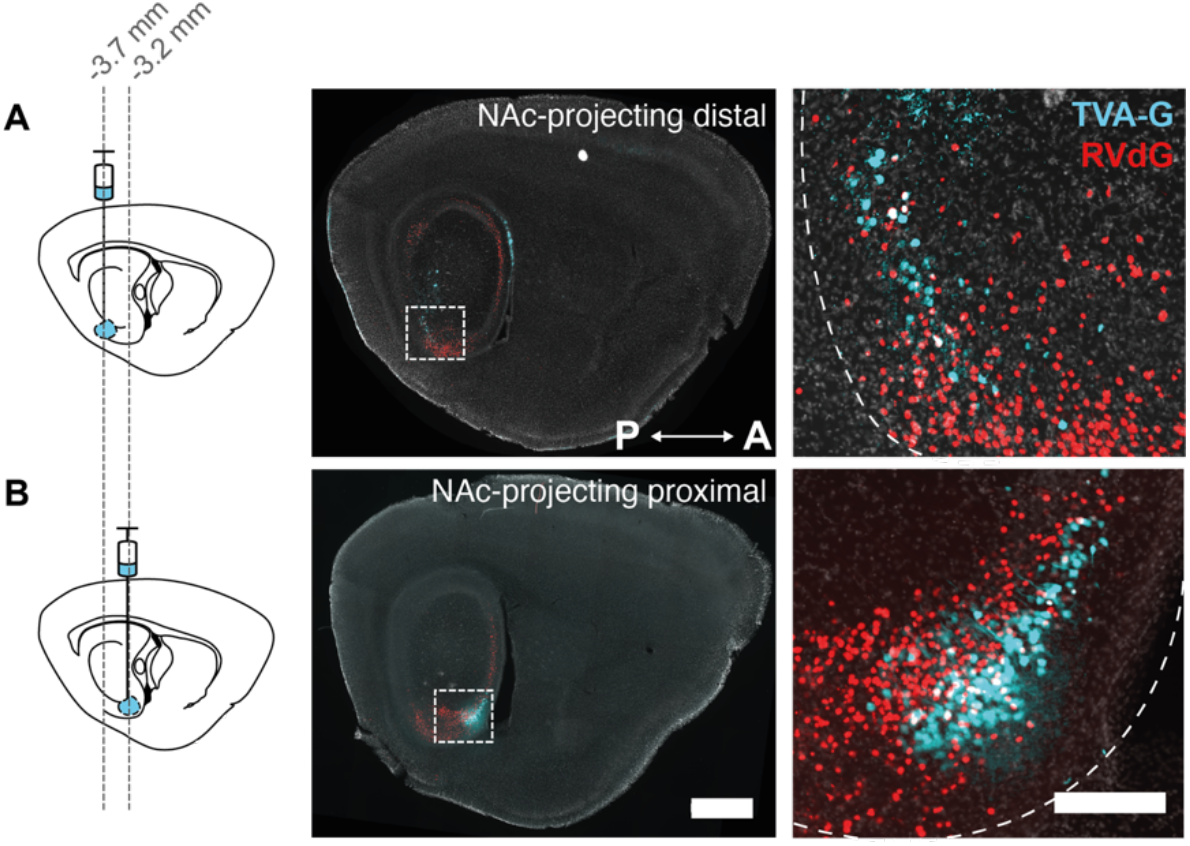
Control over starter cell location. (A – B) *Left:* Schematic to control for starter cell location. *Middle:* Sagittal sections of the hippocampus. *Right:* To control for the location of starter cells, *AAV2-retro-Cre* was injected into the target location, but *AA2/1-synP-FLEX-split-TVA-EGFP-B19G* (i.e. TVA-G) in the 1st surgery and rabies virus in the 2nd surgery were injected into either a more posterior (AP position: −3.7 mm) or anterior position (AP position: −3.2 mm). TVA-G was injected into the (A) distal position and (B) proximal position. This strategy allowed us to isolate vS^NAc^ starter cells across different proximal-distal positions (see Fig 1). Scale bars: 1000 μm (middle), 200 μm (right).

**Figure 2-figure supplement 3:**
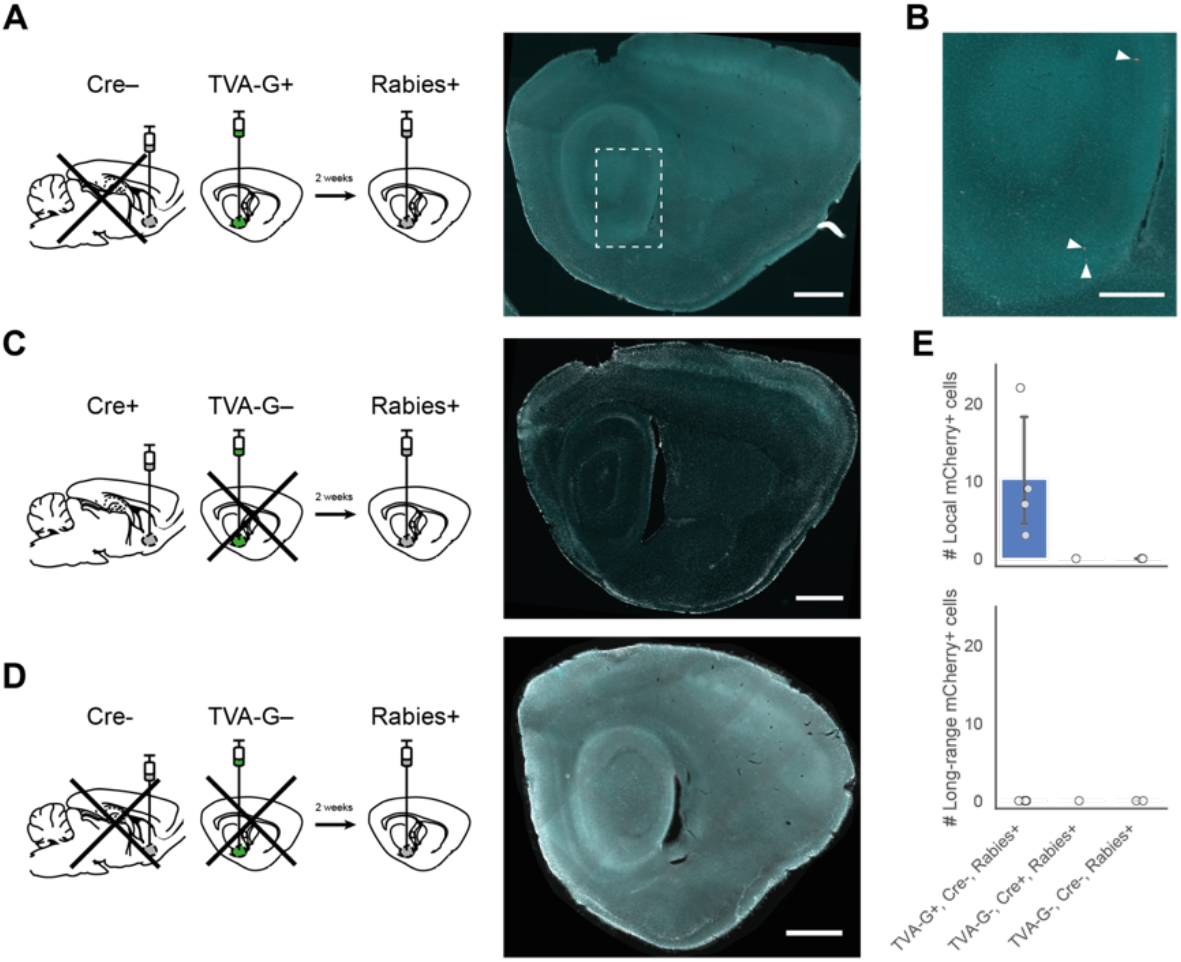
Control experiments for TRIO. (A, C, D) Schematics for control surgeries for TRIO. (A) *Left:* Controls with TVA-G injected into vS and without *AAV2-retro-Cre*, with rabies injection 2 weeks later (n = 4 brains). *Right:* Sagittal brain section. (B) Zoom-in image of boxed region in (A, middle). Arrowheads indicate sparse mCherry+ labelled rabies input cells, likely due to Cre-independent expression of TVA-G and subsequent rabies infection of starter cells. (C) No TVA-G (n = 1 brain) and (D) no TVA-G and Cre (n = 2 brains) controls. No mCherry+ cells were detected in (C) and (D). (E) Quantification of mCherry+ labelled inputs in control conditions. All leaky mCherry+ cells were detected locally within the hippocampal formation, and none were detected in long-range input regions. Bar plots indicate mean ± sem. Scale bars: 1000 μm (A,C,D, *right*), 500 μm (B, zoomed-in image).

**Figure 3-figure supplement 1:**
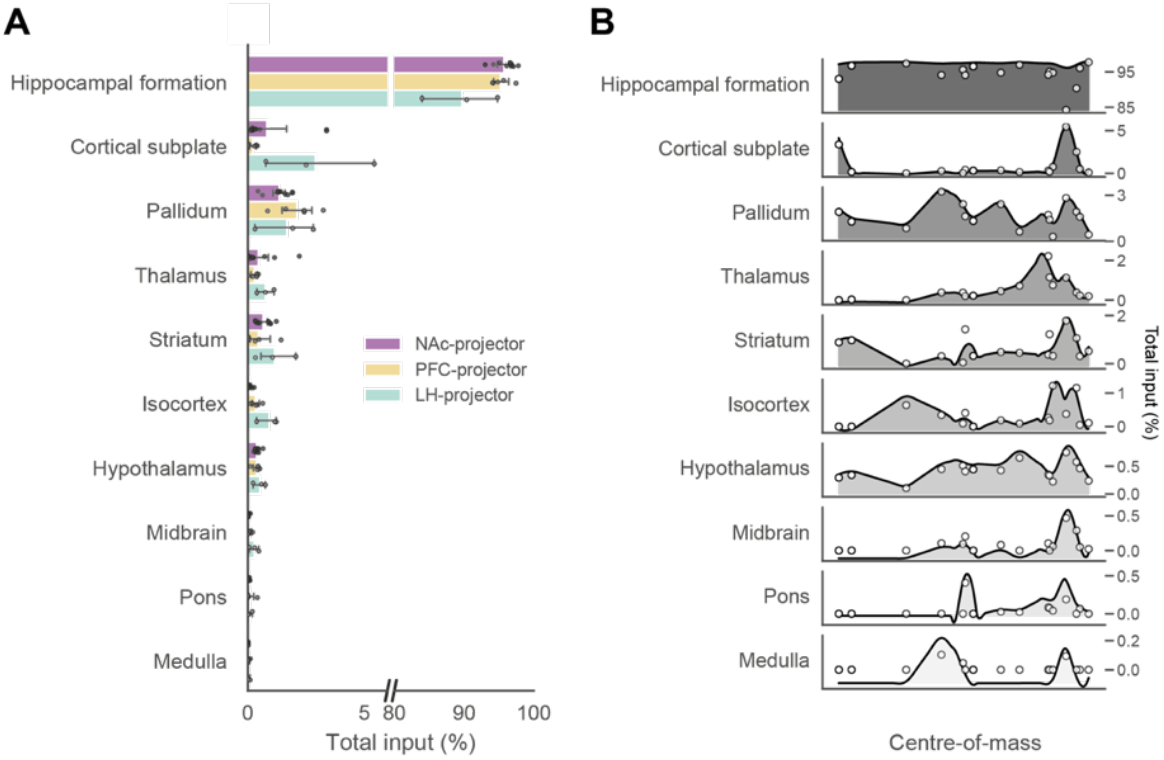
Coarse input-mapping. (A) Dataset for coarse-level inputs split according to output projection. (B) Same dataset as in A but plotted as a function of COM A-P coordinates. The shaded continuous distribution represents the smoothed input density (normalised with area under the curve = 1) as a function of COM. Bar plots indicate mean ± sem.

**Figure 3-figure supplement 2:**
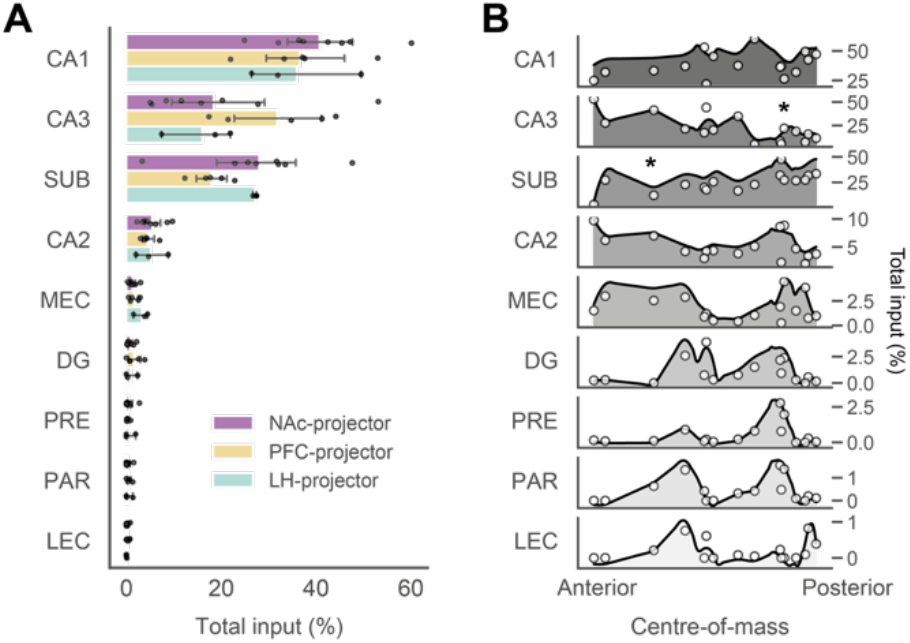
Intrahippocampal inputs. (A) Dataset of intrahippocampal inputs split according to output projection. No intrahippocampal input fractions were significantly different across projection populations after either multiple linear regression with ANOVA analysis or multiple one-way ANOVA comparisons across projection populations (see Extended Table 2). (B) Same dataset as in A but plotted as a function of COM A-P coordinates. The shaded continuous distribution represents the smoothed input density (normalised with area under the curve = 1) as a function of COM. Bar plots indicate mean ± sem. CA3 and SUB input fractions (*p < 0.05) were detected as having statistically significant correlations with COM with multiple Spearman rank correlation tests (see Extended Table 2).

**Figure 3-figure supplement 3:**
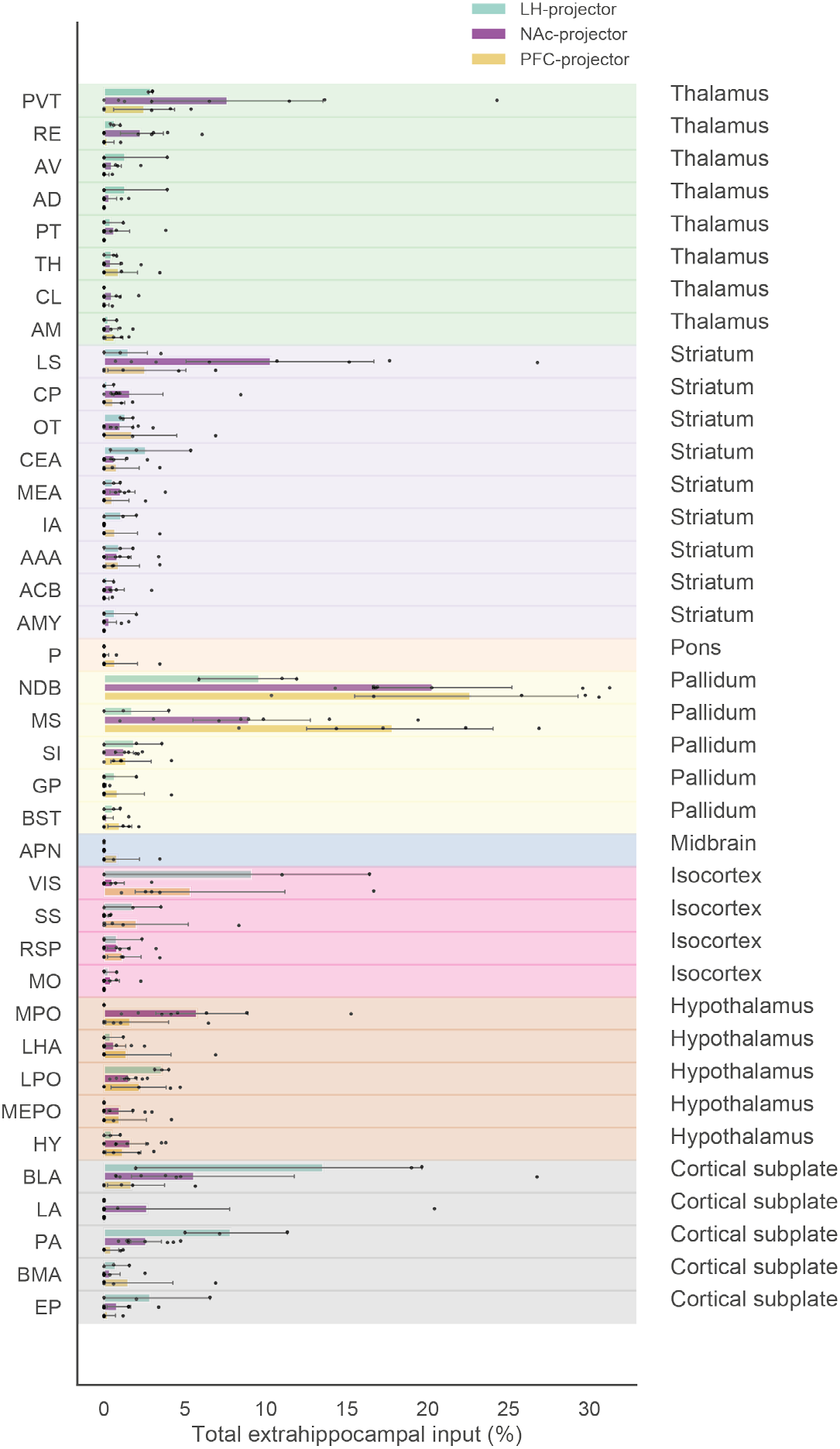
Long-range extrahippocampal inputs. Summary data for long-range inputs from multiple brain regions by each projection population. Note that the dataset was preprocessed by including only input regions that generated at least 0.05% of extrahippocampal input. Data expressed as a percentage of total input cells counted in a single brain sample. Abbreviations: see Extended Table 1. Bar plots indicate mean ± sem.

**Figure 3-figure supplement 4:**
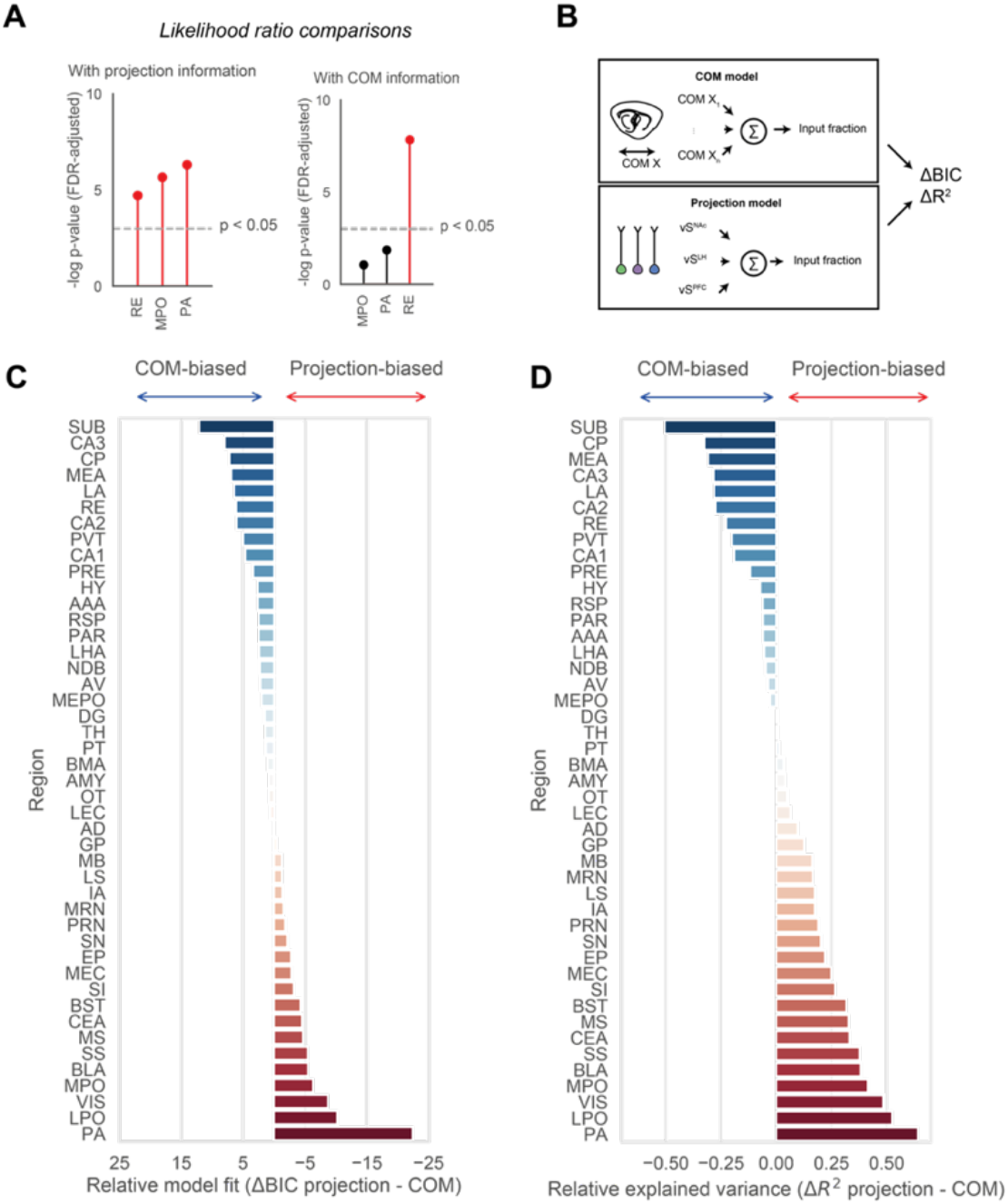
COM and projection model comparisons. (A) Single-predictor models (COM or projection only) for statistically significant model fits (MPO, RE and PA) were compared to a full model containing COM and projection identity as regressors using the likelihood ratio test. *Left:* COM-only models compared to full models. A variety of inputs - including RE and PA - are more likely to be accounted for by a model containing additional projection information with COM information. *Right:* projection-only models compared to full models. Only RE inputs are more likely to be accounted for with additional COM information as well as projection information in the full model. Note that the p-values illustrated have been adjusted for multiple comparisons of all 18 extrahippocampal regions shown in Figure 3. (B) Schematic of single-predictor linear models where COM of starter cells or projection identity were used as predictors, and the input fraction normalised to total input was used as the target variable for each brain region. The two competing models were then compared with ΔBIC and ΔR2 as measures of goodness-of-fit. (C,D) To qualitatively compare non-nested COM or projection models for each of 45 brain regions that provide input to vS, the relative likelihood (Bayesian information criterion, or BIC, score) and adjusted R2 values were calculated for each model, and the absolute difference between these values were plotted as a measure of goodness-of-fit. The more positive the ΔBIC or more negative the ΔR2, the more the input size of a given brain region is biased towards COM, while the more negative the ΔBIC or more positive the ΔR2, the more the input size is biased towards projection type.

**Figure 4-figure supplement 1:**
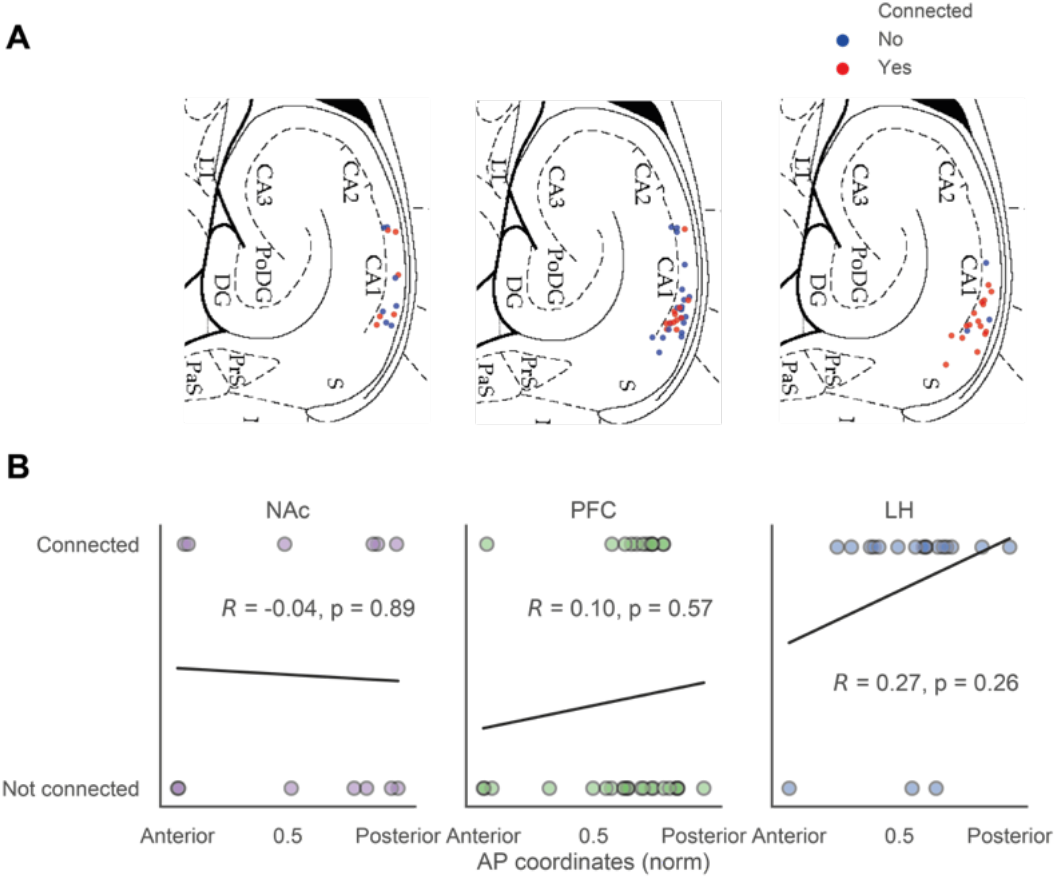
RE input connectivity by projection identity. (A) Map of RE input connectivity split by projection population. Each dot represents a patched cell, and cells were considered connected with RE input if the light-evoked photoresponse exceeds > 5 pA. (B) Input connectivity split by the projection identity of patched neurons and plotted against AP coordinates. For all projection populations, there was no statistically significant correlation between the AP coordinate and RE input connectivity of cells.

**Extended Table 1.**
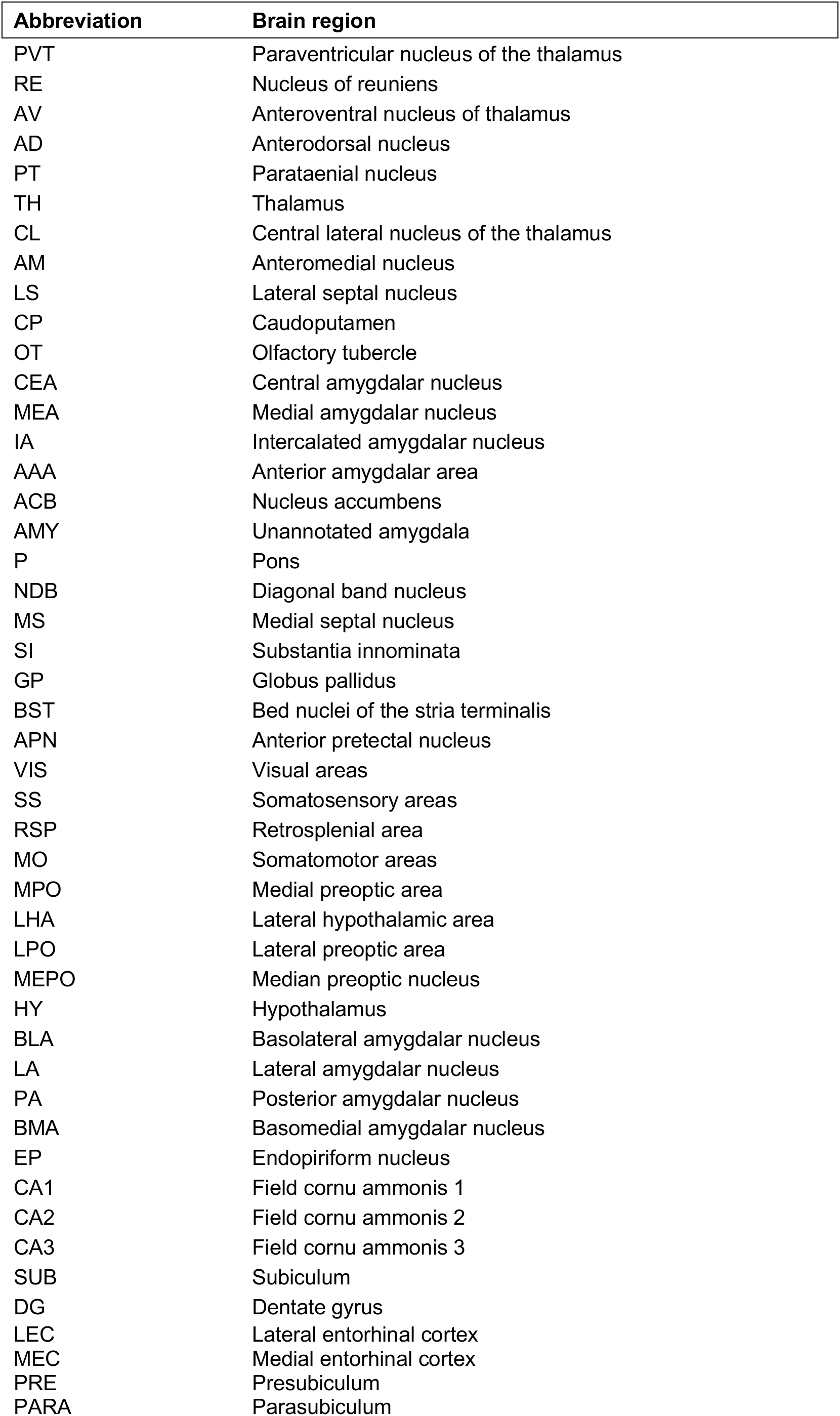

**Extended Table 2:**
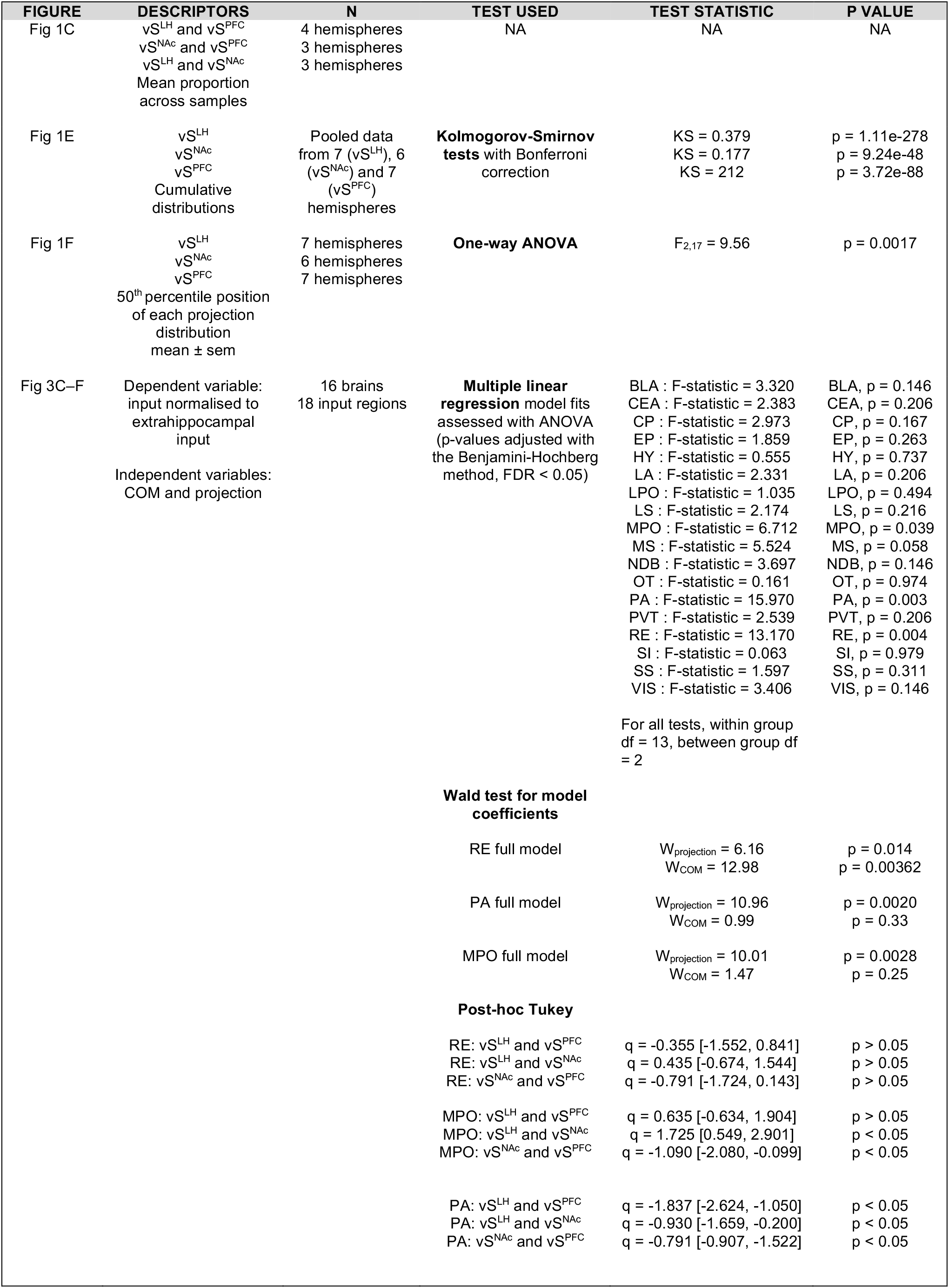
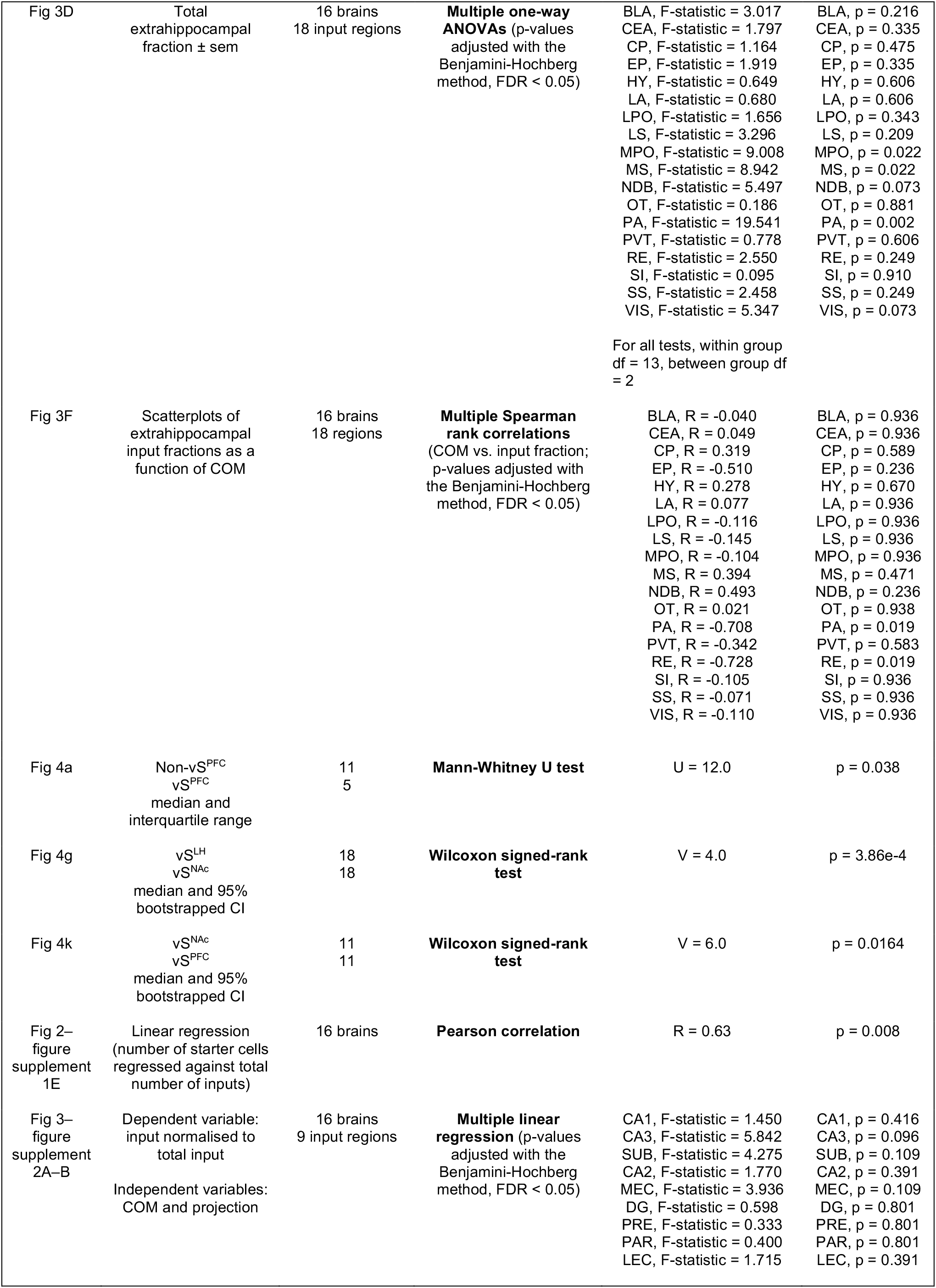
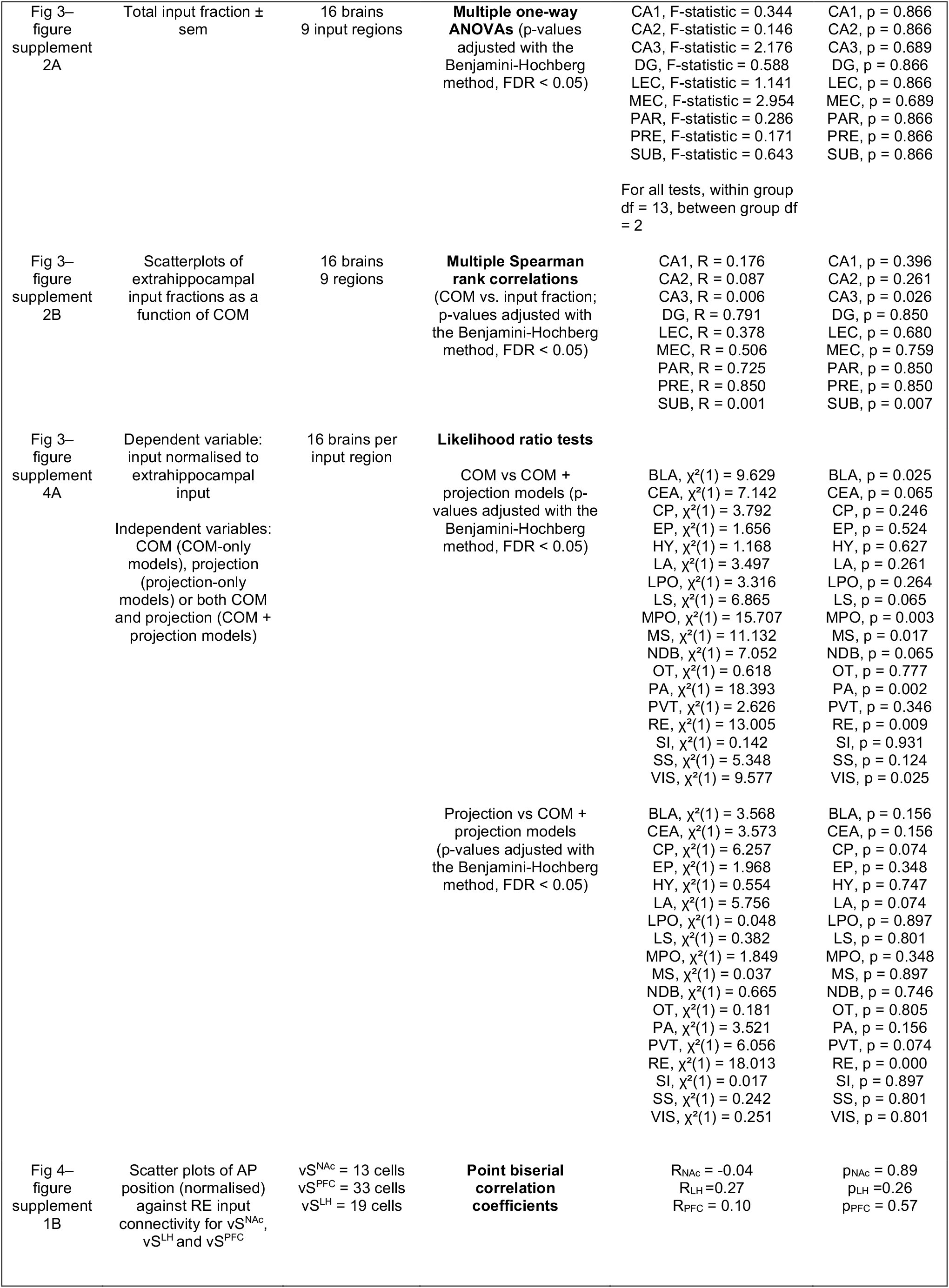
Statistical summary.

## References

Adhikari A, Topiwala MA, Gordon JA. 2010. Synchronized activity between the ventral hippocampus and the medial prefrontal cortex during anxiety. Neuron 6S:257–269. doi:10.1016/j.neuron.2009.12.002

Aggleton JP, Christiansen K. 2015. The subiculum: the heart of the extended hippocampal system. Prog Brain Res 219:65–82. doi:10.1016/bs.pbr.2015.03.003

Amaral DG, Cowan WM. 1980. Subcortical afferents to the hippocampal formation in the monkey. J Comp Neurol 189:573–591. doi:10.1002/cne.901890402

Beier KT, Gao XJ, Xie S, DeLoach KE, Malenka RC, Luo L. 2019. Topological Organization of Ventral Tegmental Area Connectivity Revealed by Viral-Genetic Dissection of Input-Output Relations. Cell Rep 26:159–167.e6. doi:10.1016/j.celrep.2018.12.040

Beier KT, Kim CK, Hoerbelt P, Hung LW, Heifets BD, DeLoach KE, Mosca TJ, Neuner S, Deisseroth K, Luo L, Malenka RC. 2017. Rabies screen reveals GPe control of cocaine-triggered plasticity. Nature 1–22. doi:10.1038/nature23888

Beier KT, Steinberg EE, DeLoach KE, Xie S, Miyamichi K, Schwarz LA, Gao XJ, Kremer EJ, Malenka RC, Luo L. 2015. Circuit Architecture of VTA Dopamine Neurons Revealed by Systematic Input-Output Mapping. Cell 162:622–634. doi:10.1016/j.cell.2015.07.015

Cembrowski MS, Bachman JL, Wang L, Sugino K, Shields BC, Spruston N. 2016. Spatial Gene-Expression Gradients Underlie Prominent Heterogeneity of CA1 Pyramidal Neurons. Neuron 89:351–368. doi:10.1016/j.neuron.2015.12.013

Cembrowski MS, Phillips MG, DiLisio SF, Shields BC, Winnubst J, Chandrashekar J, Bas E, Spruston N. 2018a. Dissociable Structural and Functional Hippocampal Outputs via Distinct Subiculum Cell Classes. Cell. doi:10.1016/j.cell.2018.03.031

Cembrowski MS, Wang L, Lemire AL, Copeland M, DiLisio SF, Clements J, Spruston N. 2018b. The subiculum is a patchwork of discrete subregions. eLife 7. doi:10.7554/eLife.37701

Cetin A, Komai S, Eliava M, Seeburg PH, Osten P. 2007. Stereotaxic gene delivery in the rodent brain. Nature Protocols 1:3166–3173. doi:10.1038/nprot.2006.450

Ciocchi S, Passecker J, Malagon-Vina H, Mikus N, Klausberger T. 2015. Selective information routing by ventral hippocampal CA1 projection neurons. Science 348:560–563. doi:10.1126/science.aaa3245

Dolleman-van der Weel MJ, Griffin AL, Ito HT, Shapiro ML, Witter MP, Vertes RP, Allen TA. 2019. The nucleus reuniens of the thalamus sits at the nexus of a hippocampus and medial prefrontal cortex circuit enabling memory and behavior. Learn Mem 26:191–205. doi:10.1101/lm.048389.118

Fürth D, Vaissiere T, Tzortzi O, Xuan Y, Martin A, Lazaridis I, Spigolon G, Fisone G, Tomer R, Deisseroth K, Carlen M, Miller CA, Rumbaugh G, Meletis K. 2018. An interactive framework for whole-brain maps at cellular resolution. Nat Neurosci 21:139–149. doi:10.1038/s41593-017-0027-7

Ito HT, Zhang S-J, Witter MP, Moser EI, Moser M-B. 2015. A prefrontal-thalamo-hippocampal circuit for goal-directed spatial navigation. Nature S22:50–55. doi:10.1038/nature14396

Jimenez JC, Su K, Goldberg AR, Luna VM, Biane JS, Ordek G, Zhou P, Ong SK, Wright MA, Zweifel L, Paninski L, Hen R, Kheirbek MA. 2018. Anxiety Cells in a Hippocampal-Hypothalamic Circuit. Neuron 97:670–683.e6. doi:10.1016/j.neuron.2018.01.016

Kim Y, Spruston N. 2011. Target-specific output patterns are predicted by the distribution of regular-spiking and bursting pyramidal neurons in the subiculum. Hippocampus 22:693–706. doi:10.1002/hipo.20931

Knierim JJ, Neunuebel JP, Deshmukh SS. 2014. Functional correlates of the lateral and medial entorhinal cortex: objects, path integration and local-global reference frames. Philos Trans R Soc Lond, B, Biol Sci 369:20130369–20130369. doi:10.1098/rstb.2013.0369

Luo L, Callaway EM, Svoboda K. 2018. Genetic Dissection of Neural Circuits: A Decade of Progress. Neuron 98:256–281. doi:10.1016/j.neuron.2018.03.040

MacAskill AF, Cassel JM, Carter AG. 2014. Cocaine exposure reorganizes cell type- and input-specific connectivity in the nucleus accumbens. Nat Neurosci 17:1198–1207. doi:10.1038/nn.3783

Masurkar AV, Srinivas KV, Brann DH, Warren R, Lowes DC, Siegelbaum SA. 2017. Medial and Lateral Entorhinal Cortex Differentially Excite Deep versus Superficial CA1 Pyramidal Neurons. Cell Rep 18:148–160. doi:10.1016/j.celrep.2016.12.012

McHenry JA, Otis JM, Rossi MA, Robinson JE, Kosyk O, Miller NW, McElligott ZA, Budygin EA, Rubinow DR, Stuber GD. 2017. Hormonal gain control of a medial preoptic area social reward circuit. Nat Neurosci 20:449–458. doi:10.1038/nn.4487

Naber PA, Witter MP. 1998. Subicular efferents are organized mostly as parallel projections: a double-labeling, retrograde-tracing study in the rat. J Comp Neurol 393:284–297.

Oh SW, Harris JA, Ng L, Winslow B, Cain N, Mihalas S, Wang Q, Lau C, Kuan L, Henry AM, Mortrud MT, Ouellette B, Nguyen TN, Sorensen SA, Slaughterbeck CR, Wakeman W, Li Y, Feng D, Ho A, Nicholas E, Hirokawa KE, Bohn P, Joines KM, Peng H, Hawrylycz MJ, Phillips JW, Hohmann JG, Wohnoutka P, Gerfen CR, Koch C, Bernard A, Dang C, Jones AR, Zeng H. 2014. A mesoscale connectome of the mouse brain. Nature S08:207–214. doi:10.1038/nature13186

Okuyama T, Kitamura T, Roy DS, Itohara S, Tonegawa S. 2016. Ventral CA1 neurons store social memory. Science 3S3:1536–1541. doi:10.1126/science.aaf7003

Ren J, Friedmann D, Xiong J, Liu CD, Ferguson BR, Weerakkody T, DeLoach KE, Ran C, Pun A, Sun Y, Weissbourd B, Neve RL, Huguenard J, Horowitz MA, Luo L. 2018. Anatomically Defined and Functionally Distinct Dorsal Raphe Serotonin Sub-systems. Cell 17S:472–487.e20. doi:10.1016/j.cell.2018.07.043

Schwarz LA, Miyamichi K, Gao XJ, Beier KT, Weissbourd B, DeLoach KE, Ren J, Ibanes S, Malenka RC, Kremer EJ, Luo L. 2015. Viral-genetic tracing of the input-output organization of a central noradrenaline circuit. Nature S24:88–92. doi:10.1038/nature14600

Soltesz I, Losonczy A. 2018. CA1 pyramidal cell diversity enabling parallel information processing in the hippocampus. Nat Rev Neurosci 1–10. doi:10.1038/s41593-018-0118-0

Strange BA, Witter MP, Lein ES, Moser EI. 2014. Functional organization of the hippocampal longitudinal axis. Nat Rev Neurosci 1S:655–669. doi:10.1038/nrn3785

van Groen T, Miettinen P, Kadish I. 2003. The entorhinal cortex of the mouse: organization of the projection to the hippocampal formation. Hippocampus 13:133–149. doi:10.1002/hipo.10037

Vertes RP. 2006. Interactions among the medial prefrontal cortex, hippocampus and midline thalamus in emotional and cognitive processing in the rat. Neuroscience 142:1–20. doi:10.1016/j.neuroscience.2006.06.027

Wyss JM, Swanson LW, Cowan WM. 1979. A study of subcortical afferents to the hippocampal formation in the rat. Neuroscience 4:463–476. doi:10.1016/0306-4522(79)90124-6

Xu W, Sudhof TC. 2013. A neural circuit for memory specificity and generalization. Science 339:1290–1295. doi:10.1126/science.1229534

O’Keefe J, Dostrovsky J. 1971. The hippocampus as a spatial map. Preliminary evidence from unit activity in the freely-moving rat. Brain Research 34:171–175. doi:10.1016/0006-8993(71)90358-1

